# Shulin packages axonemal outer dynein arms for ciliary targeting

**DOI:** 10.1101/2020.09.04.282897

**Authors:** Girish R. Mali, Ferdos Abid Ali, Clinton K. Lau, Farida Begum, Mark Skehel, Andrew P. Carter

## Abstract

The main force generators in eukaryotic cilia and flagella are axonemal outer dynein arms (ODAs). During ciliogenesis, these ∼1.8 MDa complexes are assembled in the cytoplasm and targeted to cilia via an unknown mechanism. Here we use the ciliate *Tetrahymena* to identify two novel factors (Q22YU3 and Q22MS1) which bind ODAs in the cytoplasm and are required for their delivery to cilia. We show that Q22YU3, which we name Shulin, locks the ODA motor domains into a closed conformation and inhibits motor activity. Cryo-EM reveals how Shulin stabilizes this compact form of ODAs by binding to the dynein tails. Our findings provide a molecular explanation for how newly assembled dyneins are packaged for delivery to the cilia.

---

Motile cilia play essential roles that range from setting up the left-right body axis to clearing mucus from the lungs (1). These slender cellular projections contain an axoneme built of microtubule doublets. Ciliary beating is powered by arrays of inner and outer dynein arm motors that slide adjacent doublets past each other (2). The outer dynein arms (ODAs) are the main force generators in cilia and the most frequently mutated components in human motile ciliopathies (3). ODAs are multi-subunit complexes (4), which are pre-assembled in the cytoplasm by a collection of nine dynein axonemal assembly factors and associated chaperones (4, 5). Following assembly, ODAs are targeted to cilia, where the intraflagellar transport (IFT) machinery carries them to their docking sites (6, 7). However, the mechanism of ODA delivery to the cilia and whether any additional factors are required for this process are both unknown.

To identify potential ciliary targeting factors, we purified newly-assembled ODAs from the cytoplasm of the protozoan ciliate *Tetrahymena thermophila*. We de-ciliated *Tetrahymena* to remove pre-existing axonemes and trigger new ODA synthesis (8) (**Figure 1A**). ODA complexes, containing a FLAG-tagged copy of the intermediate chain IC3, were extracted and separated from other assembly intermediates by size exclusion chromatography (SEC). Two factors co-eluted with the new fully-assembled ODAs and were identified by mass spectrometry (MS) as Q22MS1 and Q22YU3 (**Figure 1B, Figure S1A**). We also performed label-free quantitative-MS on the immunoprecipitated IC3 subunit and detected an equivalent fold enrichment of both factors relative to other ODA subunits (**Figure S1B, Table S1**). Taken together, these data suggest Q22MS1 and Q22YU3 interact tightly with ODAs in the cell body.

**Figure 1.**
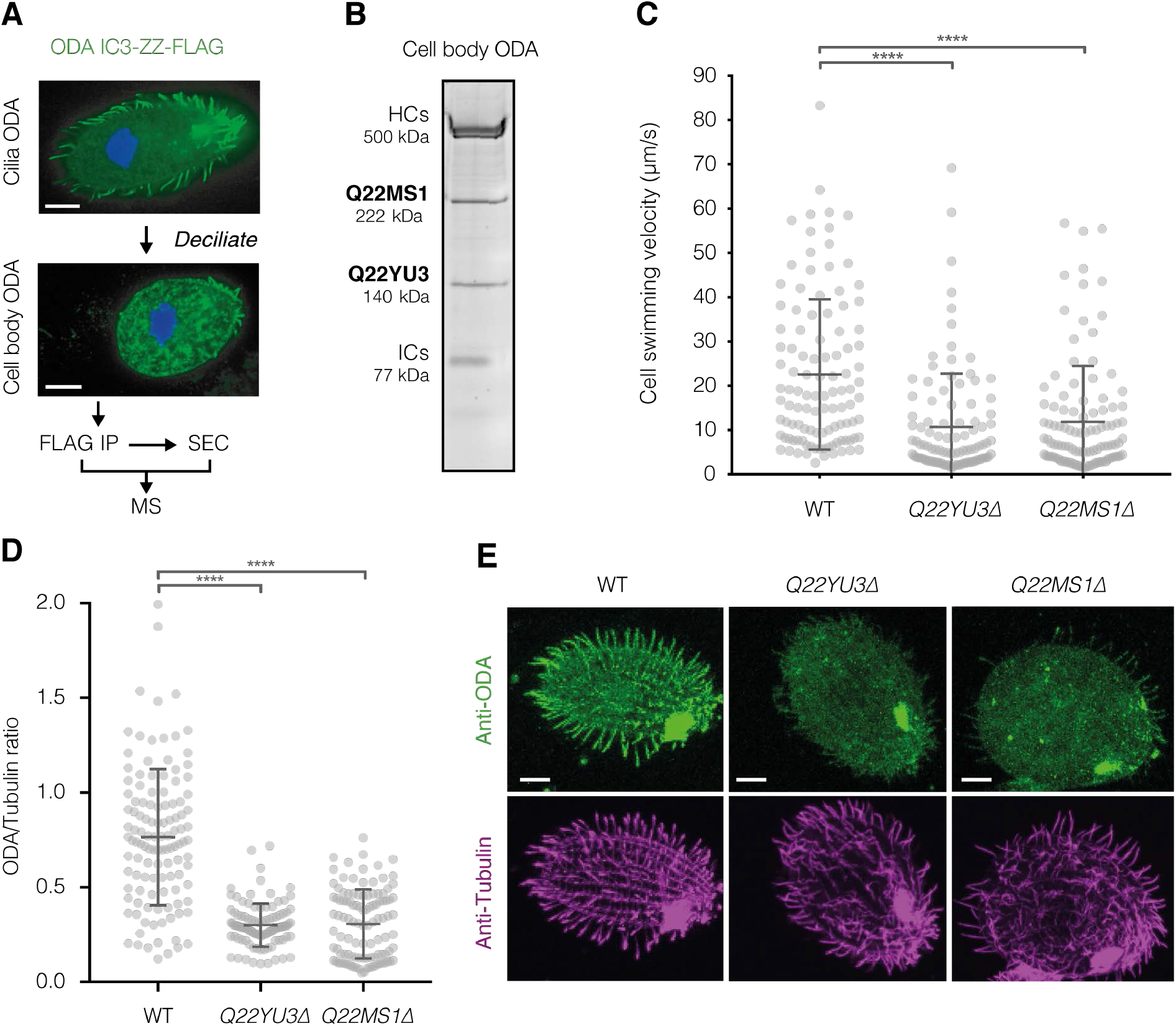
Q22MS1 and Q22YU3 deliver ODAs from cytoplasm to cilia. **(A)** Scheme used to identify novel interactors of ODAs assembled in the cell body. **(B)** SDS-PAGE of ODA purified from the cell body by IP-SEC showing co-elution with Q22MS1 and Q22YU3, HC: Heavy chains, IC: Intermediate chains. **(C)** Cell swimming velocity comparing wildtype (WT n=108) and mutant strains (*Q22YU3*Δ n=102 and *Q22MS1*Δ n=110). **(D)** Ratio of ODA/Tubulin immunofluorescence intensity along individual cilia (WT n=118, *Q22YU3*Δ n=104, *Q22MS1*Δ n=129, 3-10 cilia from 14-17 cells/genotype). **(E)** Representative cells showing immunofluorescence for ODA and tubulin (quantified in D). Scale bars:10 µm. Error bars show standard deviation; ns=not significantly different, ****p≤0.0001 (ANOVA with Tukey’s test for multiple comparisons).

These novel factors lack known functions and have not previously been linked to motile cilia. Q22YU3 shares 24% identity to human C20ORF194 (**Figure S2**) whereas Q22MS1 has a unique domain architecture and no ortholog outside of *Tetrahymena*. To investigate their roles, we generated *Tetrahymena* knockout strains for each factor (**Figure S3A**). Both strains showed an ∼2-fold decrease in swimming speed compared to wildtype (*Q22YU3*Δ: 10.6 ± 12 µm/s, *Q22MS1*Δ: 11.88 ± 12.6 µm/s, WT: 22.5 ± 16.9 µm/s; mean ± SD) suggesting defects in cilia movement (**Figure 1C**). Mutants also had decreased accumulation of food vacuoles and a higher frequency of cytokinetic defects, which are hallmarks of defective cilia motility in *Tetrahymena* (9, 10)(**Figure S3B, C**). We found the lengths and numbers of cilia in our knockouts were similar to wildtype suggesting our observations were not due to defects in ciliogenesis (**Figure S3D, E**). However, high-speed imaging showed mutant cilia beat slowly (**movie S1**), similar to a temperature-sensitive mutant with reduced ODAs in cilia (11) (**movie S2**). We therefore used immunofluorescence to test if loss of Q22YU3 and Q22MS1 affects ODA targeting to cilia. Staining with an antibody against ODAs showed marked reductions in their levels in cilia of mutant strains compared to the wildtype (**Figure 1D, E, Figure S3F**). Together, our data suggest that loss of Q22YU3 or Q22MS1 result in defective ciliary movement due to reduced delivery of ODAs to the cilia.

It has been proposed that ODAs need to be held in an inactive state during transport into the axoneme (4). We therefore tested whether Q22YU3 and Q22MS1 inhibit dynein motor activity. We expressed both factors recombinantly and assayed their effect on the microtubule gliding activity of ODAs purified from axonemes (**Figure 2A**). In the absence of Q22YU3 and Q22MS1, ODAs translocated microtubules at 1.39 ± 0.6 µm/s (mean ± SD). Microtubule gliding was severely compromised in the presence of both factors together (0.15 ± 0.15 µm/s) or with Q22YU3 alone (0.1 ± 0.15 µm/s). Q22MS1 reduced microtubule gliding velocities to a lesser extent (0.56 ± 0.76 µm/s). In contrast, addition of both factors to cytoplasmic dynein-1 did not significantly alter its microtubule gliding velocity (0.76 ± 0.47 µm/s for dynein-1 alone vs 0.69 ± 0.27 µm/ s with both factors). These data show Q22YU3 is sufficient to specifically inhibit ODA activity.

**Figure 2.**
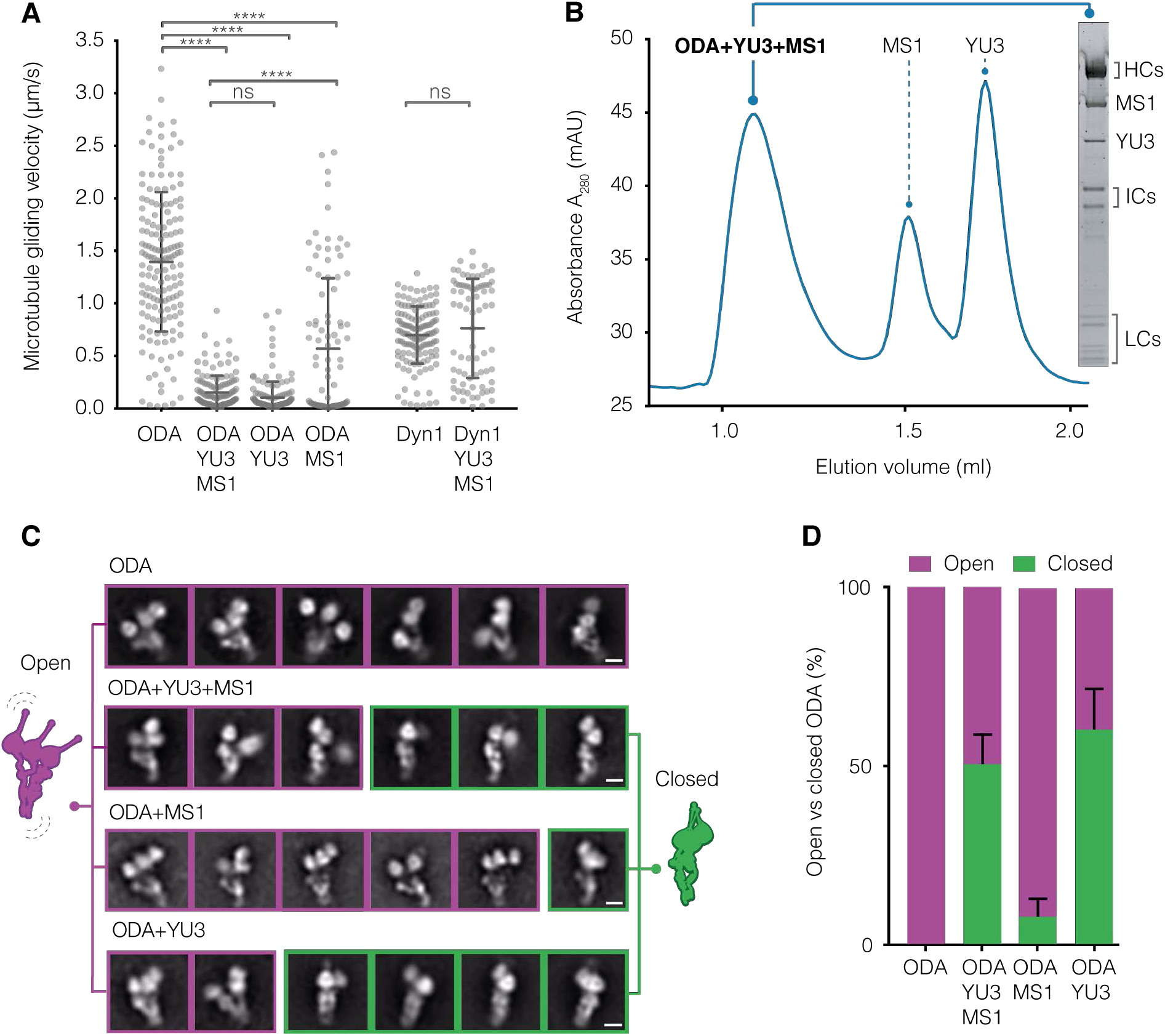
Q22YU3 binding inhibits ODAs by clustering motors. **(A)** Microtubule gliding velocities. Individual gliding events from three technical replicates/sample are plotted (ODA n=159, ODA+YU3+MS1 n=136, ODA+YU3 n=146, ODA+MS1 n=94, Dyn1 n=136, Dyn1+YU3+MS1 n=76). YU3: Q22YU3, MS1: Q22MS1, Dyn1: human cytoplasmic dynein 1. Error bars show standard deviation; ns=not significantly different, ****p≤0.0001 (ANOVA with Tukey’s test for multiple comparisons). **(B)** Axonemal-purified ODA reconstituted with recombinant Q22YU3 and Q22MS1. SDS-PAGE gel of SEC peak fraction. **(C)** Representative 2D class averages showing the distribution of open (purple) and closed (green) ODA particles from ODAs alone and reconstituted with factors. Scale bars = 10 nm. **(D)** Quantification of closed and open particles shown in (C) represented as percentage. Error bars show standard deviation.

*Tetrahymena* ODAs contain three dynein heavy chains and, when purified from axonemes, show an open bouquet conformation with the heavy chain motor domains separated (12). However, when we used negative stain electron microscopy (EM) to image ODAs purified from the cytoplasm, we noticed ∼40% of intact particles displayed a ‘closed’ conformation where the motor domains are clustered, and the tails are compacted (**Figure S4**). This closed conformation resembles a form previously observed only after cross-linking (12). To identify the factor responsible, we reconstituted ODAs extracted from axonemes with Q22MS1 and Q22YU3. Both factors form stable complexes with ODAs either together or individually (**Figure 2B, Figure S5A**). Whereas ODAs on their own were entirely in the open conformation (**Figure 2C, D**), in the presence of both factors 50 ± 8.3% (mean ± SD) of particles were closed, similar to the fraction observed for ODAs purified from the cytoplasm. ODAs bound to Q22MS1 alone were closed only 7.7 ± 5.0% of the time. In contrast, in the presence of Q22YU3 alone, 60 ± 11.4% of ODAs were closed (**Figure 2C, D, Figure S5B-F**). Collectively, our findings suggest that Q22YU3 inhibits ODAs by holding the three heavy chains together into a closed conformation. We therefore propose to name this novel protein Shulin (Sanskrit: one that holds the trident).

To elucidate how Shulin closes ODAs at a molecular level, we determined the structure of the reconstituted *Tetrahymena* ODA-Shulin complex by cryo-EM. The resolution of our overall structure is limited to 8.8 Å due to its flexibility. However, focused classification and local refinement produced a series of sub-region maps ranging from 4.3-5.9 Å in global resolution (**Figure 3A, Figure S6**). The central portion of the Shulin region map has a local resolution range of 3.2-4.2 Å (**Figure S7**) enabling de novo building of Shulin and its interactions (**Figure 3A**). In combination with MS to identify the ODA subunit composition (**Figure S8**), our maps allowed us to assign the positions of the three heavy chains (Dyh3, Dyh4, Dyh5), two essential intermediate chains (ICs: Dic2, Dic3) and 11 light chains (LCs) (**Figure 3B, Figures S6-S8**). In our structure, the motor domains are clustered in a closed conformation (**Figure 3B**) consistent with the negative stain data. At the core of the structure are the Dyh3 (α-HC) and Dyh4 (β-HC) heavy chains, which are conserved across all eukaryotes with motile axonemes (13). They form a heterodimer held together by an N-terminal dimerization do-main in an arrangement that is similar to cytoplasmic dyneins (14, 15). The third heavy chain, which is only found in ciliate and algal ODAs, is Dyh5 in *Tetrahymena* (γ-HC equivalent to the α-HC in the alga *Chlamydomonas*). Our structure shows that Dyh5 is much shorter than the other heavy chains and is anchored to Dyh4 halfway along its length. The N-terminus of Dyh5 contains a Kelch-type β-propeller domain that sits on the helical bundles of Dyh4 (**Figure 3C**). The peripheral attachment of this third heavy chain explains why its loss in *Chlamydomonas* is largely tolerated (16).

**Figure 3.**
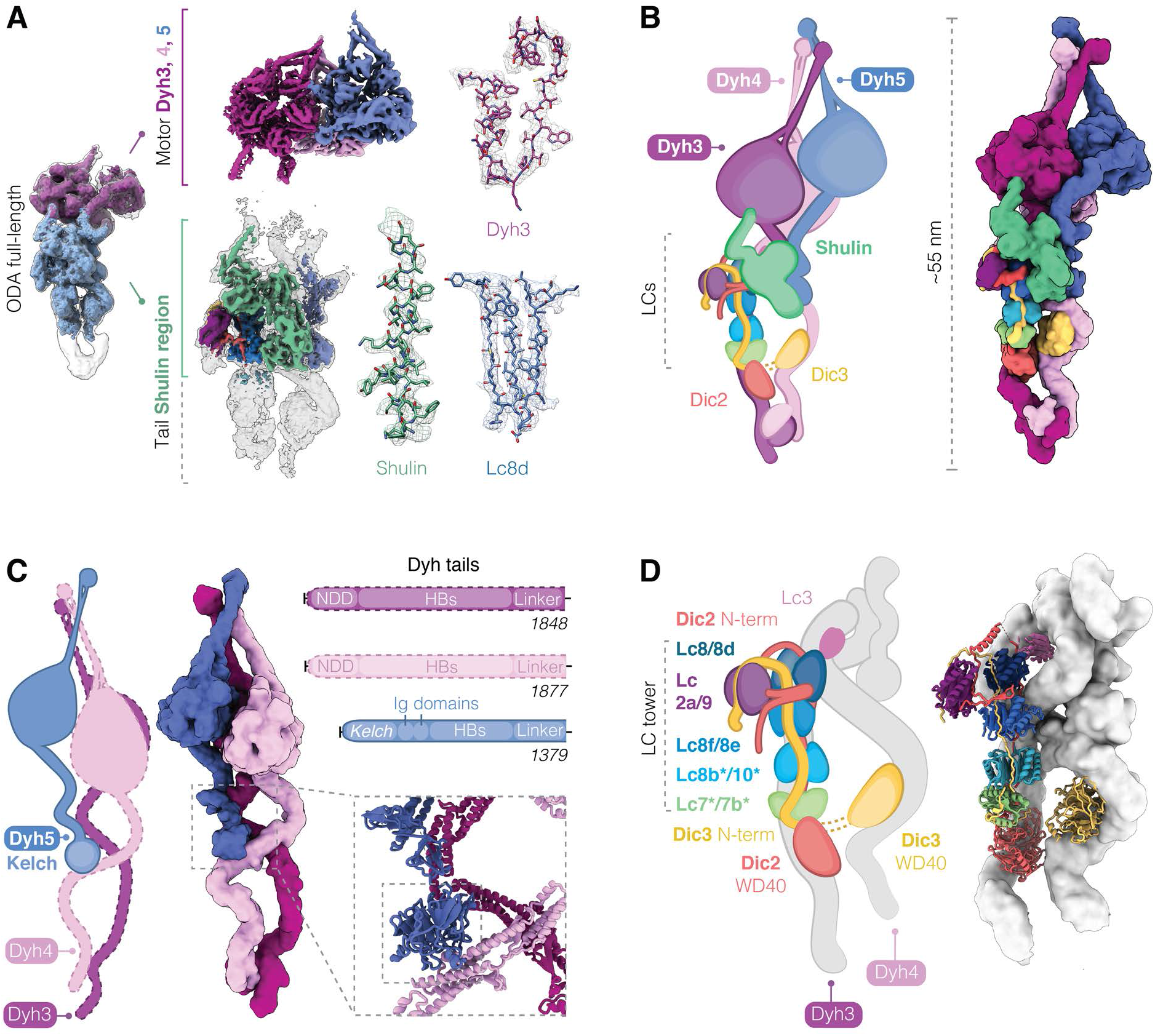
Cryo-EM reconstruction shows architecture of closed ODA. **(A)** Overview of the closed ODA bound by Shulin with head (purple) and tail (blue) maps docked in an overall map (grey). Maps obtained after masked refinements are shown for the head region containing densities for Dyh3,4 and 5 motor domains and the tail map contains a docked Shulin-region map (green). Representative cryo-EM densities are shown. **(B)** Cartoon and filtered surface representation of all modeled subunits. **(C)** Dyh5 binds Dyh4 via its N-terminal Kelch domain (inset). HB: Helical bundles, NDD: N-terminal dimerization domain. **(D)** DIC N-terminal extensions bind dimers of LCs forming a LC tower and followed by globular WD40 domains that contact Dyh3 and Dyh4. Heterodimers of Lc7/7b and Lc8b/Lc10 are tentatively assigned (*). Lc3 sits on Dyh4 and is not part of the LC tower.

In their tail regions, Dyh3 and Dyh4 wrap round the globular WD40 domains of the intermediate chains and Dyh4 also binds to a small density consistent with the thioredoxin-like Lc3 light chain (**Figure 3D, Figure S9A**). The intermediate chains have long N-terminal extensions which are held together by a tower of light chains consisting of a Lc7/Lc7b heterodimer, three dimers of Lc8 orthologs and at the end, bent over to one side, a Tctex like heterodimer of Lc2a/Lc9 (**Figure 3D, Figure S9B**). Based on side-chain density we assigned the positions of Lc8 and its orthologs: Lc8d, Lc8e and an unnamed Lc8-like protein (UniProt ID: Q22R86) which we call Lc8f. Our MS analysis showed the additional presence of Lc10 and Lc8b which we tentatively assigned to the remaining two positions. The bent arrangement of the Lc2a/Lc9 heterodimer is stabilized by the Dic2 N-terminus, which loops out from where it contacts Lc2a and wedges between Lc8d and Lc8e (**Figure S9B**).

In the motor region, all three heavy chains are locked in the pre-power stroke conformation of their catalytic cycle (17) with their force producing linker domains bent through 90°(**Figure S10A, B**). The density suggests that the coiled-coil stalks of each motor domain are angled to interact with each other close to their microtubule binding domains (**Figure 3A, B, Figure S10B**). Clustering of motor domains is further stabilized by interactions between the Dyh3 and Dyh4 linkers, between the Dyh4 AAA4 and the elbow of the Dyh3 linker and between the Dyh5 AAA3 and Dyh3 AAA4 (**Figure S10C, D**). This clustered conformation is distinct to the one ODAs adopt upon docking onto ciliary doublets where their motor domains are stacked parallel to each other and free to undergo their catalytic cycle (18). Thus, the closed conformation is an inactive state of ODAs prior to their final incorporation into cilia.

Our Shulin structure shows that it contains N-terminal domains (N1, N2) which are related to the aminopeptidase P domain of Spt16, a core component of the histone chaperone FACT (**Figure 4A**). We built the middle (M) domain *de novo*, revealing that it adopted a pleckstrin homology (PH) fold akin to the Spt16 C-terminus. Therefore, the N-terminal half of Shulin bears high structural homology to Spt16 (19). Shulin’s C-terminal domains (C1 and C2) are homologous to the bacterial GTPase YjiA (20). C1 has a Ras-like fold and C2 comprises of a five-stranded β-sheet. There is a nucleotide between C1 and C2, which plays a structural role holding the two domains together (**Figure S11A**). Shulin ends with a long C-terminal finger with a helix-loop arrangement that projects away from the core of the molecule (C3) (**Figure 4A**).

**Figure 4.**
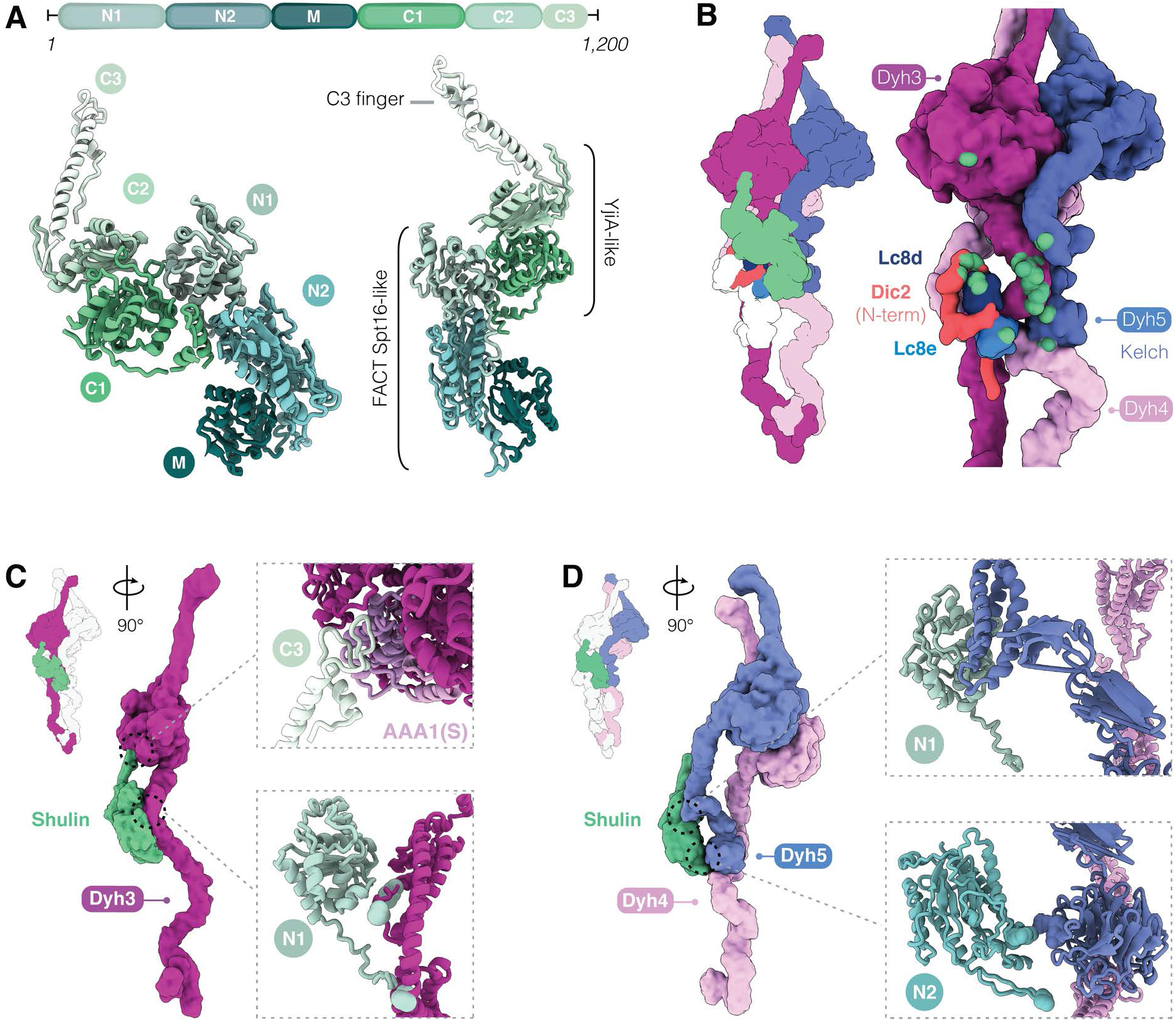
Characterization of Shulin structure and its mechanism of ODA inhibition. **(A)** Domain architecture of Shulin. N-terminal (N1, N2) and Middle (M) domains bear homology to FACT complex core subunit Spt16. C-terminal (C1, C2) domains are similar to GTPase YjiA and are followed by a C-terminal finger (C3). **(B)** Cartoon and filtered surface representation with all contacts between Shulin and ODA subunits highlighted in green spheres. **(C)** Shulin’s N1 domain contacts helical bundles proximal to the linker in Dyh3 tail. The C3 finger projects out to contact Dyh3 AAA1(S) (insets). **(D)** Shulin’s N1 domain contacts Dyh5 helical bundles and its N2 domain touches the Kelch domain. Shulin contacts Dyh4 just below Dyh5 Kelch-domain (insets).

Shulin stabilizes the closed conformation of ODAs by binding all three heavy chains and the LC tower (**Figure 4B**). It makes its most extensive interactions with Dyh3 (3216 Å^2^ surface area). Shulin’s N1 domain contacts multiple sites in the Dyh3 tail and its C3 finger binds AAA1S of the Dyh3 motor domain (**Figure 4C**). These contacts bridge the Dyh3 tail and motor holding them in a rigid conformation (**Figure S7**). Shulin can only make these connections if the motor is in its pre-power stroke conformation, suggesting it directly locks Dyh3 into its closed state (**Figure S11B**).

Shulin holds the other two motor domains in their closed conformation indirectly by stabilizing the contacts they make with Dyh3 and each other. Its N2 domain makes a small connection (313 Å^2^) to Dyh4 close to where the Dyh5 Kelch domain is docked (**Figure 4D**). This interaction holds Dyh3 and Dyh4 together and reinforces a contact between the helical bundles in their tail regions (**Figure S11B**). This, in turn, supports the previously-described connections between their motor domains (**Figure S10**). Shulin’s connection to Dyh5, via its N1 domain, is also small (292 Å^2^), but sufficient to stabilize the Dyh5 linker binding to the Dyh3 motor domain and motor-motor contacts between Dyh4 and Dyh5 (**Figure S10E**). In the light chain tower the M-domain of Shulin contacts Lc8e and its C1-domain contacts Lc8d and an -helix in the Dic2 N-terminus (**Figure S11C**). These contacts stabilize packing of the LC tower against the Dyh3 tail (**Figure S11D**). Taken together, Shulin makes contacts with multiple ODA subunits and stabilizes the interactions between them that hold the motors in a closed conformation.

Here, we identify two proteins, Q22MS1 and Shulin, which co-purify with ODAs in the cytoplasm and are required for their delivery to cilia. Q22MS1 is a 222 kDa protein containing a catalytically inactive nucleoside diphosphate kinase (NDK) domain. A homologous NDK domain is found in the recently identified *Xenopus* protein DAAP1 (21). DAAP1 has also been implicated in ODA delivery to cilia and localizes to membrane-less organelles involved in dynein assembly. Interestingly however, there is no homology between Q22MS1 and DAAP1 outside the NDK domain and the mammalian orthologs of DAAP1 lack the NDK-domain completely.

Shulin shuts down ODA motor activity, suggesting it is the proposed inhibitor (4) required during targeting of ODAs to the cilia. Unlike the cytoplasmic dyneins which are auto-inhibited (14, 15) ODAs rely on Shulin to enforce inhibition. Inactivating ODAs may be important to prevent them from escaping prematurely from cilia. Using immunostaining we found that Shulin localizes to regenerating motile cilia 30-minutes after deciliation (**Figure S12, C**). This is the stage of ciliogenesis when ODAs are being actively imported and incorporated (8). In contrast, Shulin is predominantly cytoplasmic in cells with fully assembled cilia (**Figure S12B, D**). This suggests, Shulin travels with ODAs during ciliogenesis until they reach their final axonemal location.

In addition to its inhibitory role, Shulin may directly target ODAs to cilia. A candidate for the delivery process is the small G-protein Arl3 which regulates targeting of numerous ciliary cargo (22). The human ortholog of Shulin, C20ORF194, is reported to bind the GTP-bound form of Arl3, which specifically localizes to cilia (23). Thus, Shulin may serve dual purposes of packaging and targeting ODAs to cilia.

## Materials and methods

### *Tetrahymena thermophila* strain engineering and phenotypic analyses

*Tetrahymena thermophila* wildtype *CU428* strain (Tetrahymena Stock Center) was used in this study. Cultures were maintained in SPP medium (1% bacto proteose peptone (Difco), 0.2% glucose, 0.1% yeast extract, 33 µM FeCl_3_). Transgenic lines were generated using biolistic transformation as previously described (24). For tagging IC3 polypeptide with tandem ZZ and FLAG tags, a construct bearing homology arms to the gene region upstream of the stop codon and the 3’ UTR of *DIC3* (*TTHERM_00079230*) flanking a Neomycin resistance cassette, with codon optimized sequences for the tags was used. DNA was precipitated onto 10-micron gold carriers for biolistic bombardment of the macronucleus using a Gene Gun (Bio-Rad). For disruption of *Q22YU3*/*SHULIN* (*TTHERM_00122270*) and *Q22MS1* (*TTHERM_00030520*), constructs with homology arms to the 5’ UTR and 3’ UTR of the respective genes flanking the resistance cassette were generated. Transformants were transferred to SPP medium for recovery and the promoter driving the Neomycin resistance gene was switched on by addition of cadmium chloride (1 µM). After recovery, bombarded cells were transferred into 96-well plates to isolate transformed clones. Cultures were passaged every few days into medium containing increasing concentrations of Paromomycin for phenotypic assortment of transformants. Successful generation of tagged transgenic strains was assayed by performing genomic PCRs spanning the Neo3 cassette after several generations. Only transgenic strains would amplify this resistance gene. Immunoblotting and immunofluorescence with a monoclonal FLAG antibody (Flag M2 Sigma, 1:100 dilution) further confirmed the endogenous knock-in of the epitope tag at the carboxy terminus of the IC3 polypeptide.

The knockout mutant strains tolerated up to 20-50 mg/ml of Paromomycin concentration. Genomic PCRs spanning Exons 1 to 3 of *Q22YU3* (Exon 1 forward primer: atgaatttaaattttgcatgtcttcaataag, Exon 3 reverse primer: ttatacatcatgaactgtacaatcacttgg) and *Q22MS1* (Exon 1 forward primer: atgtttggatttgaagatatccattactaacc, Exon 3 reverse primer: attagaggcttagtgaacatgtcttcgtc) confirmed disruption of both loci as only wildtype controls resulted in a robust amplicon whereas both mutants failed to generate a strong PCR product (fig. S3A). A control PCR for β-heavy chain gene was performed to verify the integrity of the genomic DNA. Additionally, immunostaining *Q22YU3*Δ/*SHULIN*Δ cells with a custom polyclonal antibody against Shulin (Eurogentec, 1:100 dilution) confirmed loss of protein as well as serving as antibody validation (**Figure S12C, D**).

For assessing ciliary defects in *Q22YU3*Δ/*SHULIN*Δ and *Q22MS1*Δ mutant strains, three main phenotypic assays were performed. Cell images or movies were acquired using a QIClick camera with QCapture software mounted onto a Leica DM IL LED microscope. Imaging was performed at room temperature. Immunofluorescence images were acquired using LSM710 confocal microscope. Videos of cells swimming close to the plane of imaging (closest to the slide) were acquired at 10 frames per second for 20-40 seconds using a 20x objective. High speed videos to visualize cilia beating were acquired by digitally magnifying on individual cells using a 100x objective at 10 frames per second. All cellular phenotyping was done using FIJI (25). Cell velocity was measured using MTrackJ plugin (26). Cell paths were manually traced, and cell velocity was calculated as a function of distance traversed in a given time frame. Food vacuoles were manually counted in phase contrast images. Cilia numbers were counted using central confocal slices in the plane of both the macro and micronucleus. Images were thresholded to segment cilia around the cell circumference for counting using FIJI. Cilia lengths were measured by drawing a line along an individual cilium and measuring its distance. Lengths of 5-15 randomly selected cilia from 14-22 cells, from replicate staining experiments were measured. Cilium lengths per cell were averaged and plotted. Cytokinesis defects were scored in images of cells stained with acetylated α-tubulin (SantaCruz, sc-23950, 1:250 dilution) to mark the cilia and demarcate the cell shape and DAPI to stain for the nuclei. Cells with more than one oral apparatus and/or macronucleus were counted for each genotype. To quantify ciliary loss of ODAs in mutants, indirect immunofluorescence was performed using a custom generated polyclonal antibody raised against the ODA holocomplex (Eurogentec, 1:150 dilution) and an acetylated α-tubulin antibody (SantaCruz, sc-23950, 1:250 dilution) to mark the ciliary axoneme. The ODA antibody was validated by the manufacturer with ELISA tests against the ODA holocomplex antigen. Additional validation of specificity was performed using a temperature sensitive *OAD1 C11* mutant strain (Tetrahymena Stock Center) with reduced ciliary ODA staining when grown at the restrictive temperature of 39°C (Attwell et al. 1992). Fluorescence intensity values along the cilium were measured using plot profile tool, averaged and plotted in GraphPad Prism7.

### Purification of ODA complexes and interactors from cell body

Large scale cultures of IC3: ZZ: 3xFLAG strains were grown to a high cell density in SPP medium typically for 72 hours. Cultures were starved in Tris-Acetate buffer for 2 hours to reduce numbers of phagocytic vacuoles containing proteases. Cells were deciliated with dibucaine hydrochloride (0.5 M). The extent of deciliation was carefully monitored by visual inspection under a stereomicroscope to ensure minimal cell lysis. Typically, within 5 minutes of adding dibucaine most cells appeared to have lost their cilia. Dibucaine concentration was diluted three-fold by adding more medium and cells were pelleted. Cell pellets were washed in Tris-Acetate buffer and lysed in Lysis Buffer (20 mM HEPES NaOH, pH 8.0, 50 mM NaCl, 1 mM EDTA, 5 mM DTT, and 10% glycerol supplemented with 0.1% Triton X-100, 0.2% IGEPAL CA-630, 1 mM PMSF, 5 µM proteasome inhibitor MG-132, and 3x Complete protease inhibitor tablets (Roche). Lysates were clarified by ultracentrifugation at 70,000 rpm in a Ti70 rotor (Beckman) and flown multiple times over FLAG-M2 affinity beads (Sigma) which were pre-equilibrated in lysis buffer and packed in a gravity flow column. The beads were washed for at least 5 column volumes times in Wash buffer (20 mM HEPES, pH 7.4, 50 mM NaCl, 1 mM MgCl_2_, 1 mM TCEP, 10% glycerol and 0.1% IGEPAL). IC3-ZZ-3xFLAG containing complexes were eluted into 5 fractions by sequentially flowing 5 bed volumes of elution buffer containing FLAG peptide. Efficiency of elution was assessed by running SDS-PAGE gels and staining with Instant Blue (Expedeon). Elution of desired ODA complexes was deemed successful upon detection of high molecular weight bands corresponding to dynein heavy chains. Eluted fractions containing the highest concentration of complexes were further fractionated over a Superose 6 increase 3.2/300 size exclusion column (GE Healthcare) in GF50 gel filtration buffer (25 mM HEPES pH 7.4, 50 mM NaCl, 1 mM MgCl_2_, 1 mM DTT, 0.1 mM ATP). FLAG eluates and peak fractions from gel filtration run were analysed by mass spectrometry.

### Purification of ODA from cilia axonemes

Axonemal ODAs were purified as previously described (12). Large scale wildtype *Tetrahymena* cultures were deciliated using dibucaine (as above). Deciliated cell pellets were discarded and cilia in the supernatant were centrifuged at 13,500 x *g* at room temperature. Cilia pellets were washed in Cilia Isolation Buffer (CIB: 20 mM HEPES pH7.4, 100 mM NaCl, 4 mM MgCl_2_, 0.1 mM EDTA) and centrifuged at 600 x *g* several times to remove cell bodies and mucus. A final high-speed spin at 13500 x *g* at 4°C yielded a pure cilia pellet with a white fluffy appearance. Cilia were demembranated by resuspending the pellet in CIB containing 0.25% Triton-X detergent, freshly added protease inhibitors, 1 mM DTT and 200 mM PMSF followed by a 30-minute incubation on ice. Demembranated cilia axonemes were isolated by centrifugation at 17000 x g at 4°C. Axonemes were washed in CIB buffer to remove residual detergent and repelleted. Axoneme pellets were resuspended in high salt buffer (20 mM HEPES pH7.4, 600 mM NaCl, 4 mM MgCl_2_, 0.1 mM EDTA, 1 mM DTT, 0.1 mM ATP, 200 mM PMSF) and incubated for 30 minutes on ice to isolate dynein arms. The dynein containing high salt extract was loaded onto 6 identical 5-25% sucrose density gradients made in CIB and ODA arms were separated from other axonemal complexes over 16 hours by centrifugation at 33,000 rpm in an SW40 rotor at 4°C. The following day, sucrose gradients were manually fractionated into 500 µl fractions. Every alternate fraction was resolved on an SDS-PAGE gel and stained with instant blue. ODA isolation was deemed successful upon detection of characteristic high molecular weight bands corresponding to ODA heavy chain polypeptides over several of the denser fractions towards the bottom of the gradient. Sucrose gradient fractions containing ODA complexes were further purified over a MonoQ 5/50 anion exchange column (GE Healthcare) to separate out other axonemal dynein species. ODA complexes eluting at ∼300 mM salt off the MonoQ column were verified for intactness and presence of subunits by a further gel filtration step, negative stain electron microscopy and mass spectrometry analyses. ODA complexes purified as above were used in all biochemical reconstitution experiments and subsequent cryo-EM studies.

### Insect cell expression and purification of Q22MS1 and Q22YU3/Shulin

Gene sequences coding for *Tetrahymena thermophila* Q22MS1 (*TTHERM_00030520*) and Q22YU3 (*TTHERM_00122270*; C20ORF194-like) were codon-optimized and synthesized (Epoch) for expression in *Spodoptera frugiperda* derived *Sf* 9 cells. Codon optimized sequences were sub-cloned into pACEBac1-derived vectors containing C-terminal 2xStrep tag. The following constructs were generated pACEBac1-Q22MS1-Psc-2ŒStrep and pACEBac1-Q22YU3-Psc-2ŒStrep. Baculoviruses for individual expression of Q22MS1 and Q22YU3/Shulin were prepared using the insect cell-baculovirus system. Cells expressing recombinant proteins were harvested 48 hours after infection and lysed in 50 ml cell lysis buffer (20 mM Hepes-NaOH pH 7.2, 100 mM NaCl, 2 mM MgAc, 1 mM EDTA, 10% (v/v) glycerol, 1 mM DTT). Cells were mechanically lysed in a 40 ml Dounce-type homogenizer (Wheaton) using 15-25 strokes. Lysates were clarified by ultracentrifugation at maximum speed in a Ti70 rotor (503,000 x *g*) for 45 min, 4°C (Beckman Coulter). Clarified lysates were poured several times over 0.5-1 ml Streptactin resin (IBA) which was applied into a gravity flow column and pre-equilibrated in lysis buffer. The resin was washed for 20 column volumes to remove non-specifically bound contaminants. Recombinant proteins were eluted off the resin in five fractions over sequential incubations in an elution buffer (lysis buffer containing 3 mM D-desthiobiotin). Eluates were resolved on an SDS-PAGE and stained with Instant Blue to assess the purity of the recombinant proteins. Recombinant proteins eluting off the Streptactin beads were further cleaned over gel filtration using GF150 (25 mM HEPES pH 7.4, 150 mM NaCl, 1mM MgCl_2_, 1 mM DTT, 0.1 mM ATP) buffer. Proteins were snap frozen in liquid nitrogen and stored at -80°C for use in all biochemical reconstitutions which were performed at 4°C.

### Reconstitution of ODA with Shulin and/or Q22MS1

ODA complexes (∼0.5-1 mg/ml) purified over a MonoQ column were mixed with 10-25x molar excesses of purified Q22YU3/Shulin, Q22MS1 (∼1-1.5 mg/ml) or both. Complexes formed by overnight dialysis into 50 mM or 150 mM NaCl buffer were used for negative stain EM analyses and complexes reconstituted by dialysis into 150 mM salt buffer were used for cryo-EM grid freezing and analyses. Complex formation was assessed by fractionating dialysates over a Superose 6 increase 3.2/300 size exclusion column (GE) in GF50 (25 mM HEPES NaOH pH7.4, 50 mM NaCl, 1mM MgCl_2_, 1 mM DTT, 0.1 mM ATP) or GF150 (25 mM HEPES NaOH pH 7.4, 150 mM NaCl, 1mM MgCl_2_, 1 mM DTT, 0.1 mM ATP) buffer. Fractions spanning the entire column volume were resolved on an SDS-PAGE gel and stained with Coomassie or SYPRO Ruby gel stain (Bio-Rad). The primary peak consisted of ODA subunits with bound factors confirming successful reconstitution. Secondary peaks contained molar excesses of individual proteins unbound to ODAs.

### Microtubule gliding assays and quantification

Gliding assays were performed as previously described (27). Microtubules were polymerized at 37°C using Alexa-647 and unlabeled tubulin at 3 µm and 11 µm concentration respectively in polymerization mix (BRB80: 80mM potassium PIPES pH6.8, 1mM MgCl_2_ and 1mM EGTA with 10mM GTP). Polymerized microtubules were stabilized with 2 µM Taxol in BRB80 (Sigma T1912). 10-20 µl of freshly prepared or freshly thawed ODA at ∼100 µg/ml concentration was applied to a flow chamber and allowed to adhere to glass for 2 min at room temperature. Flow chamber was washed in buffer (50 mM KAc, 10 mM HEPES pH 7.4, 4 mM MgAc, 1 mM EGTA) containing 1% BSA. Taxol stabilized microtubules were flowed in with a motility mix (20 mM HEPES pH 7.4, 5 mM MgSO_4_, 1 mM DTT, 1 mM EGTA) and allowed to bind to motors. Finally, motility mix containing 1mM ATP was flowed in prior to TIRF imaging. Imaging was performed at room temperature using a Nikon microscope with a 100x oil-immersion objective (Nikon, 1.49 NA Oil, APO TIRF).

The imaging system used the 100 mW 641 nm (Coherent Cube) laser. Images were acquired with a back illuminated EMCCD camera (iXonEM+ DU897E, Andor, UK) controlled with µManager software. Imaging was performed with 100 ms exposures taken at 2 s intervals, one pixel = 0.16 × 0.16 µm with a pixel size of 160 nm. For testing effect of factors, ODAs were pre-incubated on ice for 30 minutes with Q22YU3/Shulin, Q22MS1 or both in 10-25x molar excess and these complexes were applied to the flow chamber as above. These molar ratios resulted in successful reconstitutions as above and enforced closure of ODAs (by Shulin) and were therefore also used in gliding assays. As control, full-length human cytoplasmic dynein-1 (Schlager et al. 2014) was used at 100 µg/ml with or without both factors. Microtubule gliding velocities for each condition tested were calculated by manually tracking the leading edge of moving microtubules using the FIJI plugin MTrackJ. The average velocity for the track of each microtubule was used to calculate the average velocity for the entire population of microtubules recorded. Experiments were performed in triplicate technical repeats.

### Negative stain electron microscopy analysis

Negative stain microscopy analyses were performed on ODAs from axonemes, ODAs from cell bodies and ODA complexes reconstituted with Q22YU3/Shulin, Q22MS1 or both. In each case, 3 µl of sample at ∼0.05-0.1 mg/ml concentration were applied for 1 minute to freshly glow-discharged 400 mesh copper grids coated with a continuous carbon support layer (Agar Scientific) followed by application of 2% uranyl acetate for a minute and air-dried after wicking away excess liquid. For statistical analysis of open versus closed ODA bouquets, triplicate datasets per condition were collected manually on a FEI Spirit T12 microscope (equipped with Gatan 2K Œ 2K CCD (model 984) operated at 120 kV with a 1-1.5 second exposure and a pixel size of 3.64-3.5 Å/pix. For initial structural analysis of cell body ODA complexes, large datasets were collected using EPU on a FEI F20 microscope operated at 200 kV with 1s exposure; 3.4 Å/pix. A typical dose of ∼20 e^1^ per Åand a range of defoci between 0.5-1.5 µm were used.

### Statistical analysis of open and closed ODA conformations

To quantify effect of factors on ODA structure, ODAs alone, ODAs reconstituted with Q22YU3/Shulin, Q22MS1 or with both factors were each freshly prepared in triplicates and applied to negative stain grids. Micrographs were collected manually from random regions of grids and processed using RELION 3 (28). CTF was estimated using GCTF (29). A small subset of manually picked particles from each dataset yielded class averages that were used to autopick particles for that dataset.

Several rounds of 2D classification were performed to remove ambiguous particles representing ODAs which had fallen apart or could not be assigned into closed or open classes. Only intact three headed ODAs were further sub-classified. From each of the three datasets per sample, a total of 13299 (ODA), 9118 (ODA+Q22YU3+Q22MS1), 9966 (ODA+Q22YU3) and 3610 (ODA+Q22MS1) intact ODA particles were used. These particles were sub-classified and assigned into open versus closed conformations. Open conformation refers to ODAs with heads (motor domains) far apart and tails open in a V-shape kinked to one side. In contrast, a closed state is characterized by a tightly clustered appearance of heads and a compact tail straight in line with the heads. Further rounds of sub-classification were done, and raw particles were visually inspected to assess that particles were being correctly assigned into an open or closed class according to their conformation. All four datasets were analysed in triplicates (triplicate datasets acquired from triplicate reconstitutions per sample) and for each set mean frequency of open versus closed and standard deviations were calculated.

For cell body ODAs, a large dataset of 279,106 particles acquired from a cell body purification of ODAs was sub-classified as above. Non-ODA particles such as ribosomes, other cellular complexes and broken ODAs were classed in an ambiguous class. Intact ODA particles were sub-classified as above until closed and open classes were clearly distinguishable (**Figure S4**).

### Cryo-EM grid preparation

ODA were reconstituted with Q22YU3/Shulin using 1:10-1:25 molar ratios (ODA:Shulin; ODA at ∼0.5-1 mg/ml and Shulin at ∼1-1.5 mg/ml) as described above. Reconstituted complexes were purified over a Superose 6 increase 3.2/300 size exclusion column SEC in GF150 buffer and immediately crosslinked in 0.025% Glutaraldehyde (Sigma-Aldrich) for 30 minutes on ice followed by quenching with 1 mM Tris-HCl (pH 7.4). This mild crosslinking with 0.025% glutaraldehyde minimized complex dissociation during grid freezing. To assess that crosslinking did not cause gross artefacts to reconstituted complexes, crosslinked complexes were negatively stained and had an appearance indistinguishable to non-crosslinked complexes freshly applied to EM grids.

ODA:Shulin complexes were applied at a concentration of ∼0.1-0.2 mg/ml to graphene oxide (GO) grids. GO grids were prepared a day prior to freezing as previously described (30). Briefly, gold 300 mesh Quantifoil R2/2 holey carbon grids (Quantifoil Micro Tools) were glow discharged using an Edwards Sputter Coater 305B. Graphene oxide (GO) dispersion (Sigma-Aldrich; 2 mg/mL in H2O) was diluted ten-fold with ddH2O to a final concentration of 0.2 mg/ml and subsequently spun down at 600 x *g* for ∼15 sec to remove large aggregates of GO flakes. Three microliters of GO flake solution from the top was applied to grids. After incubation for one minute with graphene oxide dispersion, the GO solution was removed by blotting briefly with Whatman No.1 filter paper and washed by absorbing 20 µl ddH2O onto the GO coated side twice and once on the back side of the grid with blotting steps in between. Grids were air-dried and used the next day for cryo-EM grid freezing without further glow discharging as GO grids were already hydrophilic. Freezing was performed at 4°C with 100% humidity. 3 µl of sample were applied to the GO side of the grids. After a wait time of 45 seconds, excess liquid was blotted away for 2-2.5 seconds with Whatman filter papers pre-equilibrated in the humidity chamber. Grids were immediately plunge-frozen into liquid ethane using a Vitrobot IV (ThermoFisher Scientific). Grids were transferred into grid-boxes and stored in liquid nitrogen for future screening and data collection.

### Cryo-EM data collection and initial processing of whole ODA molecule

Electron micrograph movies were recorded using a Titan Krios (Thermo Fisher Scientific) equipped with an energy filtered K3 detector (Gatan) at 81,000x magnification in EFTEM mode. Six datasets were collected at the LMB using Serial EM at a pixel size of 1.11 Å/pixel, 300 kV, 66 frames, 2.6 s exposure, ∼52 e^-^/Å^2^. A script was used to collect data in a 3×3 hole pattern, 3 images/ hole, using beam-tilt to speed up data collection. A further dataset was collected at the Electron Bio-Imaging Centre (eBIC), Diamond Light Source, UK using EPU v2.7 at 0.53 Å/pixel, 300 kV, 54 frames, 3 s exposure, 54 e^-^/Å^2^ (**Table S2**). Aberration-free image shift (AFIS) collection was used to speed up data collection (4 images/hole).

### Cryo-EM image processing

All image processing was performed using RELION-3.1 and software wrapped within (28). Inter-frame motion in each movie was corrected using RELIONs own implementation of motion correction as described above, using 5×5 patches and a B-factor of 150 Å applied to the micrographs (31). Defocus parameters were estimated on non-dose weighted micrographs either using GCTF v1.18 (29) or CTFFIND4 v4.1.13 (32). For each individual dataset, particles were picked on non-dose weighted micrographs using Gautomatch v0.56 using permissive picking parameters (cc_cutoff=0.1, 400 Å diameter) and projections from an initial structure of full-length ODA. This initial model was obtained from a preliminary round of processing of the eBIC and first two LMB datasets where the phi-dynein structure was used as a reference (EMD-3705).

Particles were extracted from dose weighted micrographs with bin 4 parameters (768-pixel box size rescaled to 192, yielding a final pixel size of 4.24-4.44 Å/pixel). 2D classification was subsequently performed with 75-100 classes, T=4, 750 Å circular mask, limiting resolution E-step to 15 Å and ignoring CTFs until first peak. A further 2-3 rounds of 2D classification without alignment (50-70 classes) was performed to remove graphene oxide layer artifacts, ice contamination and aggregated particles refractory to averaging. Particles from 2D classes showing projections of recognizable ODA views were selected and joined from all datasets (1,300,000 particles).

The combined particles were subjected to global auto-refinement with a loose mask around the whole ODA and an initial model of the whole molecule filtered to 50 Å giving a 15 Å structure. Considerable flexibility was observed in this overall structure (hereafter referred to as overall-1), particularly the Dyh4 and Dyh5 motors and the lower tail section. To resolve the tail and motors of ODA separately, we employed a combination of focused classification, masked refinement and signal subtraction as outlined in **Figure S6**. All masks used were created using volumes of sub-regions generated in Chimera (UCSF) using ‘volume eraser’ or ‘color zone’ (33). These sub volumes were low pass filtered to 15 Å with a soft edge and binary map extension (both 6-8 pixels).

### Processing of full-length and Dyh4/Dyh5 motors

To improve upon the resolution of the full-length ODA structure, the overall-1 refinement data.star was used as input for masked 3D classification without alignment (5 classes, T=4). Particles from the class showing the most complete density (evidence for Dyh4 and Dyh5 at lower threshold levels) were selected for masked global 3D refinement, yielding a 9.7 Åstructure. At this point particles were re-extracted to their bin by 2 and unbinned parameters in parallel: 384-pixel box size at 2.22 Å/pixel and 768-pixel box size at 1.11 Å/pixel, respectively. These particles were used for local refinements, yielding full-length ODA structures that resolved to 8.9 Å (bin by 2) (overall-2) and 8.8 Å (unbinned) (overall-3; EMD-11576). A mask was applied to the tail of the overall-3 map and a local refinement was performed resulting in a 6.7 Å map of the entire ODA tail (EMD-11577). The overall-2 map provided higher signal to noise ratio for the flexible Dyh4 and Dyh5 motor domains and was thus used to resolve these regions through signal subtraction and focused refinements. To this end, particle subtraction was performed by centering the subtracted images on a mask encompassing both Dyh4 and Dyh5 motors and re-boxing to 256 pixels (2.22 Å/pixel). Subtracted particles were subjected to a local refinement, giving a an overall 12.9 Å structure of Dyh4-Dyh5 motors. This reconstruction was used as the basis for signal subtraction of Dyh4 and Dyh5 individually. Each motor was subsequently locally refined (Dyh4, 10.5 Å and Dyh5, 11.8 Å with no post processing) and 3D classified without alignment (5 classes, T=50). A final local refinement with particles from the best 3D classes (selection criteria: ordered motor, presence of stalk and limited noise) was performed, resolving to 5.0 Å (57,761 particles) and 5.6 Å (49,756 particles) for Dyh4 (EMD-11582) and Dyh5 (EMD-11583, EMD-11584), respectively.

### Processing of the tail and Shulin region

To get high resolution information on the Shulin region, a mask for the full tail was first generated based on the overall-1 structure. Using this mask, a round of 3D classification without alignment was performed on bin by 4 particles (8 classes, T=8) with the data.star from overall-1 as input. Particles from the class containing clear density for the Shulin finger, mid- and low tail region were selected for global refinement, resolving to 9.0 Å. This was subsequently used as an input for 3D classification without alignment with a mask focusing on the Shulin region (8 classes, T=100). Six classes containing clear density for the Shulin region were selected for a local refinement that produced an 8.9 Å reconstruction. At this point, 123,484 particles were unbinned and re-boxed to a smaller 384-pixel box size (1.11 Å/pixel). A more specific mask of Shulin was created encompassing its core N-terminal and C-terminal domains, the C3 finger as well as contacts to the uppermost LC tower and portions of the contacting Dyh3-5 tails. Masked local refinement of this region produced a 4.8 Å structure. Finally, the Shulin region refinement parameters were used for a round of 3D classification without alignment to sort out remaining heterogeneity (5 classes, T=50). 43,338 particles from two overlapping classes were combined for two separate masked local refinements of the Shulin region: Shulin region and the DYH3 tail contact (4.6 Å; EMD-11580), and Shulin region excluding C3 finger (4.3 Å; EMD-11579). A larger mask of the tail was also applied to the earlier 123,484 subset of particles to get an overview of the lower tail region (refined to 5.9 Å after signal subtraction, re-centering and masked 3D classification; EMD-11578).

### Processing of the Dyh3 region

The overall-1 map indicates that Dyh3 is rigid relative to the two other motors, enabling direct masked 3D classification without alignment of this motor and region of the upper tail (based on overall-1 data.star, 8 classes, T=10). Selection of particles from the best class and global 3D refinement resulted in a 9.0 Å structure, at which point particles were unbinned and reboxed (512-pixel box size). A tighter Dyh3 motor only mask was generated and used for a round of local 3D refinement, producing a 4.8 Å structure. This was used as the basis for 3D classification without alignment to separate out conformational heterogeneity (5 classes, T=50), resulting in only one class showing complete density. 49,397 particles from this class were selected for a final round of local 3D refinement and re-centering in the box, yielding a 4.4 Å resolution structure (Dyh3 region; EMD-11581).

### Model building and refinement

Homology models were generated for ODA subunits using PHYRE2 (34), using the sequences of chains found in mass spectrometry. These were supplemented by homology models from sequences for all ODA light chain subunits. Initially models were fit into density using rigid body fitting, followed by jiggle fitting in Coot (35). For the dynein heavy chain, density in the motor domains allowed for the distinction between Dyh3 and Dyh4. For the LC tower, side-chain density allowed us to confidently assign four of the LC8-like light chains (W7XJB1_Lc8, Q24CE5_Lc8d, Q24DI9_Lc8e, Q22R86 (named Lc8f), and the Tctex-like light chains (A4VEB3_Lc9 and Q1HGH8_Lc2a). The roadblock light-chains Lc7 and Lc7b were tentatively assigned based on their differing C-terminal sequence lengths. The final two LC8-like chains identified in the MS data were assigned to the remaining positions based on their N-terminal sequence lengths. Models in higher resolution density were manually refined into maps using Coot (EMD-11579, EMD-11581), rebuilding regions when necessary. Side-chain resolution density also allowed us to distinguish the N-termini of the two intermediate chains, Dic2 and Dic3. The orientation of the two N-termini allowed us to distinguish the IC WD40 domains, which were modeled. We could also assign the registry of Dyh3 and Dyh4 in the tail regions that contacts Shulin (EMD-11580).

For Shulin, PHYRE2 initially generated two models. The N-terminus residues (21-500 aligned with c5ce6A) were predicted to adopt a fold similar to Spt16 from the FACT complex, whilst the residues (726-1104 aligned with c1nijA) were predicted to fold similar to the GTPase YjiA. These two domains were fit into EMD-11579 in Coot (35). We then rebuilt sections of both domains into our map and built the middle domain of Shulin de novo. The C terminal finger was fit into lower resolution density, with the loops that contact the Dyh3 motor domain built using Dyh3 region map EMD-11581. Density for nucleotide between the C1 and C2 domains was fitted with a GTP analogue from PDB 2HF8, changed to GTP and manually refined in Coot. Once all the subunits were modeled or built, the structure was split, with subunits refined against the highest resolution maps using PHENIX (36) and REFMAC5 (37) (table S3). Regions were refined until their model validation statistics, calculated using PHENIX, no longer improved. In other regions, homology models were docked into density (EMD-11577, EMD-11578, EMD-11582, EMD-11583, EMD-11584) then refined using PHENIX, including secondary structure restraints. For all three motor domains, microtubule binding domains were tentatively placed based on the angles and registries of the stalks. All regions were then reassembled into one model, with boundaries refined using PHENIX. For the overall model, all side chains were removed for deposition.

### Mass spectrometry

Protein identification by mass-spectrometry was done either using in-gel or in-solution tryptic digestion. Three separate types of MS analyses were performed. 1) For identifying novel interactors of cell body ODAs, IC3-ZZ-FLAG pulldowns were performed on deciliated cells (as described above) in quadruplicates. Eluates were run on SDS-PAGE gels, Coomassie stained and excised gel slices were analysed by mass spectrometry. 2) For identifying proteins in the cell body ODA peak fraction following pulldowns and gel filtration, samples were resolved on SDS-PAGE and stained with SYPRO ruby protein gel stain (Bio-Rad). Polyacrylamide gel slices containing the bands for ODA holocomplex and bound factors were prepared for mass spectrometric analysis. 3) For precisely identifying subunit composition of ODAs in cilia, IP/MS experiments as above were performed on the isolated ciliary fraction of the IC3-ZZ-FLAG strains. Additionally, MonoQ fractions containing ODAs purified from cilia of wildtype strains (as described above) were tryptically digested in-solution for mass spectrometric analysis. Both these latter mass spectrometry experiments identified the same subunit composition for ODAs.

For polyacrylamide gel slices (1-2 mm) containing the purified proteins were prepared for mass spectrometric analysis by manual in situ enzymatic digestion. Briefly, the excised protein gel pieces were placed in a well of a 96-well microtitre plate and destained with 50% v/v acetonitrile and 50 mM ammonium bicarbonate, reduced with 10 mM DTT, and alkylated with 55 mM iodoacetamide. After alkylation, proteins were digested with 6 ng/µL Trypsin (Promega, UK) overnight at 37 řC. The resulting peptides were extracted in 2% v/v formic acid, 2% v/v acetonitrile. Protein samples in solution were reduced with 10 mM DTT and alkylated with 50 mM iodoacetamide. Following alkylation, the proteins were digested with trypsin (Promega, UK) at an enzyme-to-substrate ratio of 1:100, for 1 hour at room temperature and then further digested overnight at 37 řC following a subsequent addition of trypsin at a ratio of 1:20. The digests were analysed by nano-scale capillary LC-MS/MS using a Ultimate U3000 HPLC (ThermoScientific Dionex, San Jose, USA) to deliver a flow of approximately 300 nL/min. A C18 Acclaim PepMap100 5 µm, 100 µm x 20 mm nanoViper (ThermoScientific Dionex, San Jose, USA), trapped the peptides prior to separation on a C18 Acclaim PepMap100 3 µm, 75 µm x 150 mm nanoViper (ThermoScientific Dionex, San Jose, USA). Peptides were eluted with a gradient of acetonitrile. The analytical column outlet was directly interfaced via a modified nano-flow electrospray ionization source, with a hybrid dual pressure linear ion trap mass spectrometer (Orbitrap Velos, ThermoScientific, San Jose, USA). Data dependent analysis was carried out, using a resolution of 30,000 for the full MS spectrum, followed by ten MS/MS spectra in the linear ion trap. MS spectra were collected over a m/z range of 3002000. MS/MS scans were collected using a threshold energy of 35 for collision induced dissociation. LC-MS/MS data were then searched against an in-house protein sequence database, containing Swiss-Prot and the protein constructs specific to the experiment, using the Mascot search engine program (Matrix Science, UK) (38).

Database search parameters were set with a precursor tolerance of 5 ppm and a fragment ion mass tolerance of 0.8 Da. Two missed enzyme cleavages were allowed and variable modifications for oxidized methionine, carbamidomethyl cysteine, pyroglutamic acid, phosphorylated serine, threonine, tyrosine, tertbutyloxycarbonyl-lysine, norbornene-lysine and prop-2-yn-1-yloxycarbonyl-lysine were included. MS/MS data were validated using the Scaffold program (Proteome Software Inc., USA) (39). All data were additionally interrogated manually. For quantitative analysis of replicate runs, NSAF (Normalized Spectral Abundance Factor) values for each protein hit were used as a proxy of protein abundance (40). NSAF values were used to calculate significance and fold-changes (i.e. consistent enrichment of a given protein in test sample over control sample).

### Bioinformatics

Predicted homology models were generated using Phyre2 (34). For ortholog searches and sequence alignments PSI-BLAST and align tools embedded in UniProt, NCBI, ENSEMBL and Tetrahymena Genome Databases were used. Sequence alignments were visualized using ESPript (41).

### Image processing and structure representation tools

FIJI (25) used for all phenotyping and gliding assays. RELION-3.1 (42) was used for all EM data processing.Cryo-EM maps wereresampledusingEMDA(https://www2.mrc-lmb.cam.ac.uk/groups/murshudov/content/emda/emda.html). Chimera (33) and ChimeraX (43) used for model fitting and density map figure making. Manuscript formatting for Biorxiv used a modified LaTeX template from the Henriques Lab (https://www.overleaf.com/latex/templates/henriqueslab-biorxiv-template/nyprsybwffws#.Wp8hF1Cnx-E)

### Statistical tests and data representation

Usage of one-way ANOVA with Tukey’s test for multiple comparisons is indicated wherever applied. High speed videos of cells to visualize cilia movement were composited using the kapwing online tool. All graphical representations were generated using GraphPad Prism 7.

## Data and materials availability

Atomic coordinates and cryo-EM maps have been deposited in the Protein Data Bank under accession code 6ZYW, 6ZYX, 6ZYY and in the Electron Microscopy Data Bank under accession codes EMD-11576, EMD-11577, EMD-11578, EMD-11579, EMD-11580, EMD-11581, EMD-11582, EMD-11583, EMD-11584.

## Author contributions

G. R. M. discovered Shulin, performed all cell biological, biochemical and negative stain EM analyses, optimized and prepared ODA-Shulin cryo-EM samples. C. K. L collected all cryo-EM datasets. G. R. M., F. A. A. and C. K. L. determined the cryo-EM structure. A. P. C., C. K. L. and F. A. A. built and refined models. M. S. and F. B. performed mass spectrometry. A. P. C. and G. R. M. conceived the project, oversaw its implementation, and with F. A. A. prepared figures and wrote the manuscript. All authors contributed to aspects of manuscript writing and editing.

## ACKNOWLEDGEMENTS

We thank B. Santhanam for help with comparative genomic analyses, J. Grimmett and T. Darling for data storage and high-performance computing support, MRC LMB EM Facility (G. Sharov, G. Cannone) and Diamond (eBIC: proposal bi23268) for microscopy data acquisition support, K. Nguyen and V. Chandrasekaran for microscopy time, F. Coscia for GO grid assistance, S. Bullock and members of the Carter lab for comments on the manuscript, C. Stone for figure scripting and S. Utekar for suggesting the name Shulin.

---

**Figure S1.**
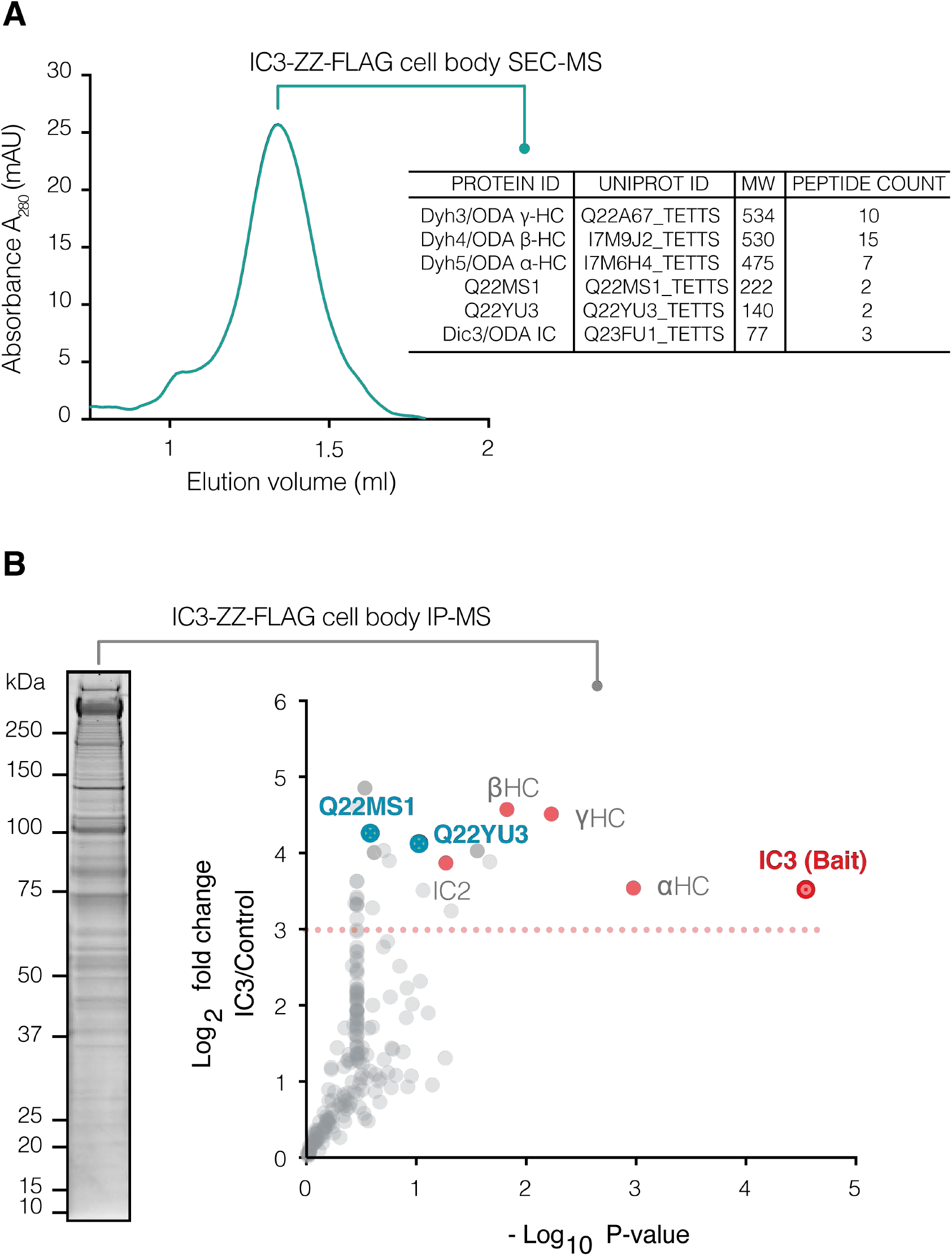
Purification and identification of novel interactors of cell body ODAs. **(A)** SEC trace showing major peak of cell body ODA complex. Co-eluting proteins were identified by MS. Molecular weight (MW) in kDa and peptide counts per protein are shown in the table. **(B)** SDS-PAGE gel showing IC3-ZZ-FLAG immunoprecipitates from deciliated *Tetrahymena* cell bodies. Protein bands were identified by mass spectrometry. Plot summarizes top hits enriched >3 fold (dotted line) in IC3 (bait protein encircled in red) over untagged control immunoprecipitates from 4 replicate experiments. Q22MS1 and Q22YU3 are highlighted (blue circles) as novel interactors.

**Figure S2.**
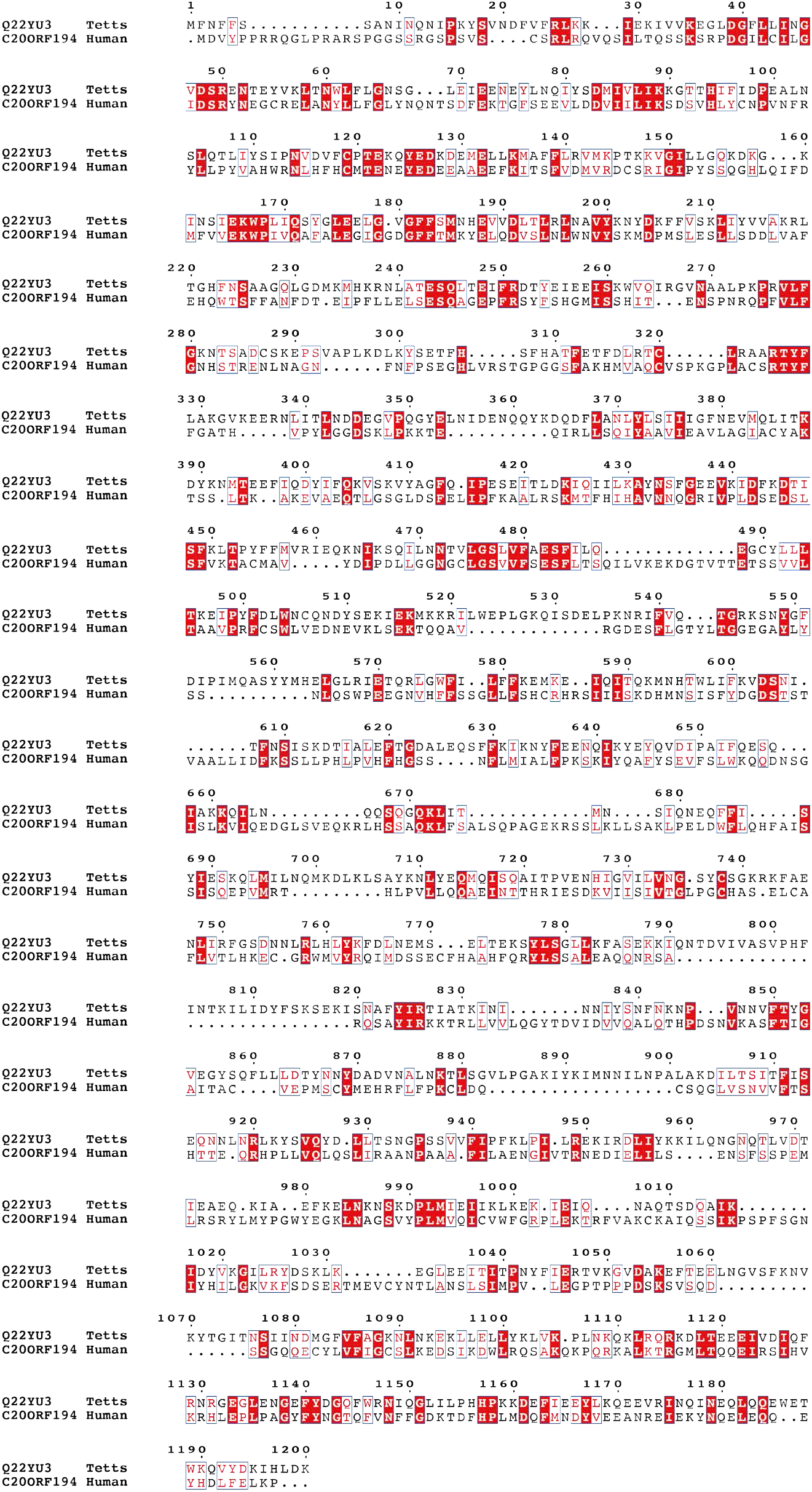
Alignment between Q22YU3 and human C20ORF194. Sequence alignment between *Tetrahymena* Q22YU3 (Q22YU3_TETTS: UNIPROT ID: Q22YU3) and human C20ORF194 (CT194_HUMAN, UNIPROT ID: Q5TEA3) shows 24% identity highlighting evolutionary conservation. Conserved residues are highlighted in red and similar residues are colored in red with blue boxes around.

**Figure S3.**
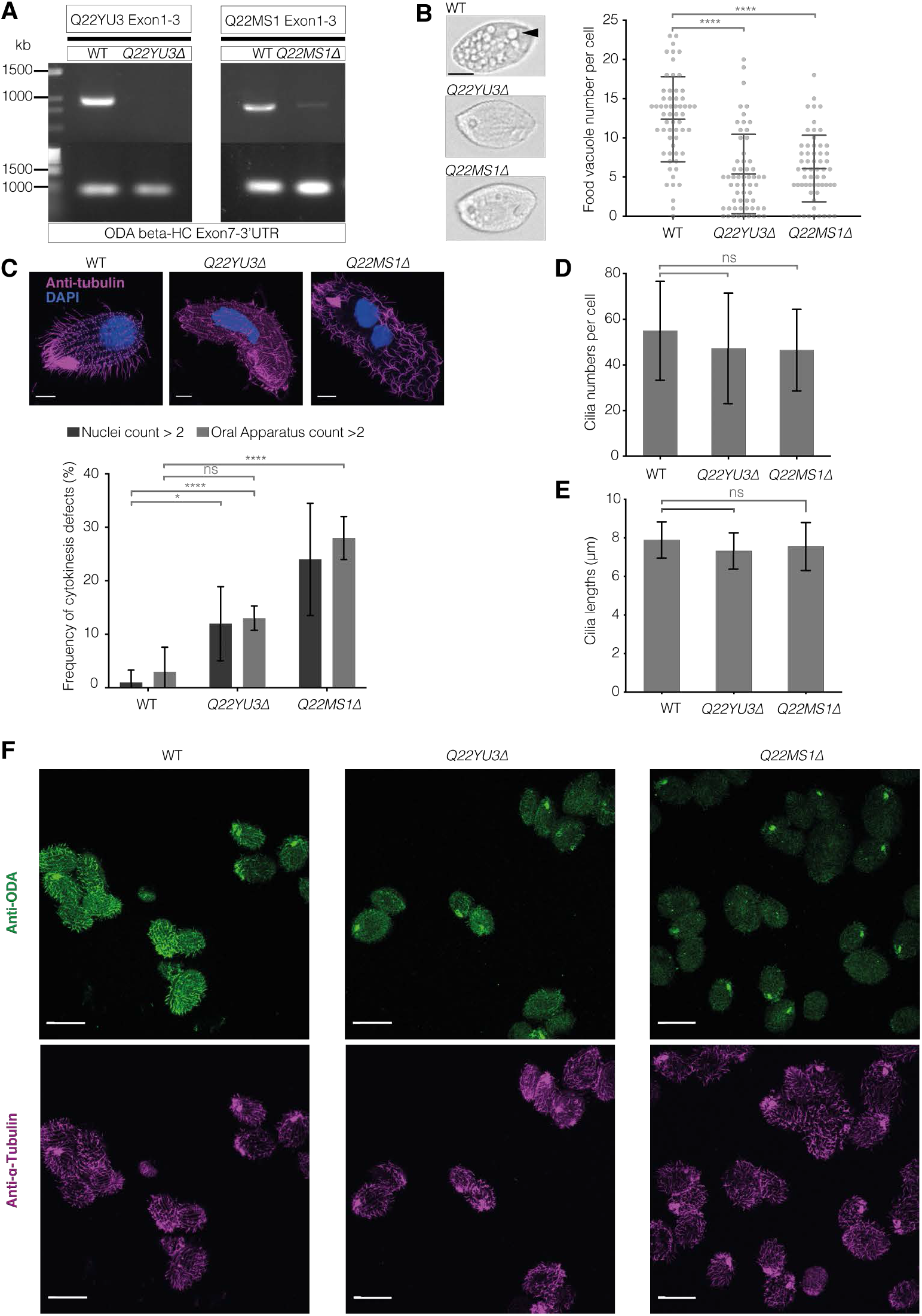
Phenotypic characterization of *Q22YU3*Δ and *Q22MS1*Δ mutant strains. **(A)** Genomic PCRs show amplicons generated using primers spanning exons 1-3 of *Q22YU3* and *Q22MS1* in wildtype (WT) and mutant strains. Control PCRs for *DYH4* (ODA β-HC) spanning exon 7 and 3’ untranslated region (UTR) of the gene are shown. (B) Cell images and graph show mutant cells have a near absence or highly reduced number of food vacuoles (arrowhead) compared to wildtype cells. N=59 cells per genotype. Scale bar = 10 µm. (C) Immunofluorescence images of wildtype and mutant cells with quantification of cytokinesis defects show *Q22YU3*Δ and *Q22MS1*Δ cells have a higher frequency of binucleated cells and multiple oral apparatuses compared to wildtype cells. N=25 cells/genotype. Two-way ANOVA was used to calculate p-values; ns=0.07, *p≤0.04, ****p≤0.0001. Scale bar = 10 µm. (D) Cilia numbers show mutants have similar numbers of cilia to wildtype cells (WT n=31, *Q22YU3*Δ n=25, *Q22MS1*Δ n=24). (E) Average cilia lengths are similar between mutant and wildtype cells (WT n=18 cells, 146 cilia; *Q22YU3*Δ n=16 cells, 147 cilia; *Q22MS1*Δ n=22 cells, 238 cilia). (F) A larger field of view showing representative cells immunostained for ODA and acetylated α-tubulin (related to Figure 1E). Scale bar = 50 µm. Error bars where used show standard deviation. One-way ANOVA was used unless otherwise indicated to calculate p-values; ns=not significantly different, ****p≤0.0001.

**Figure S4.**
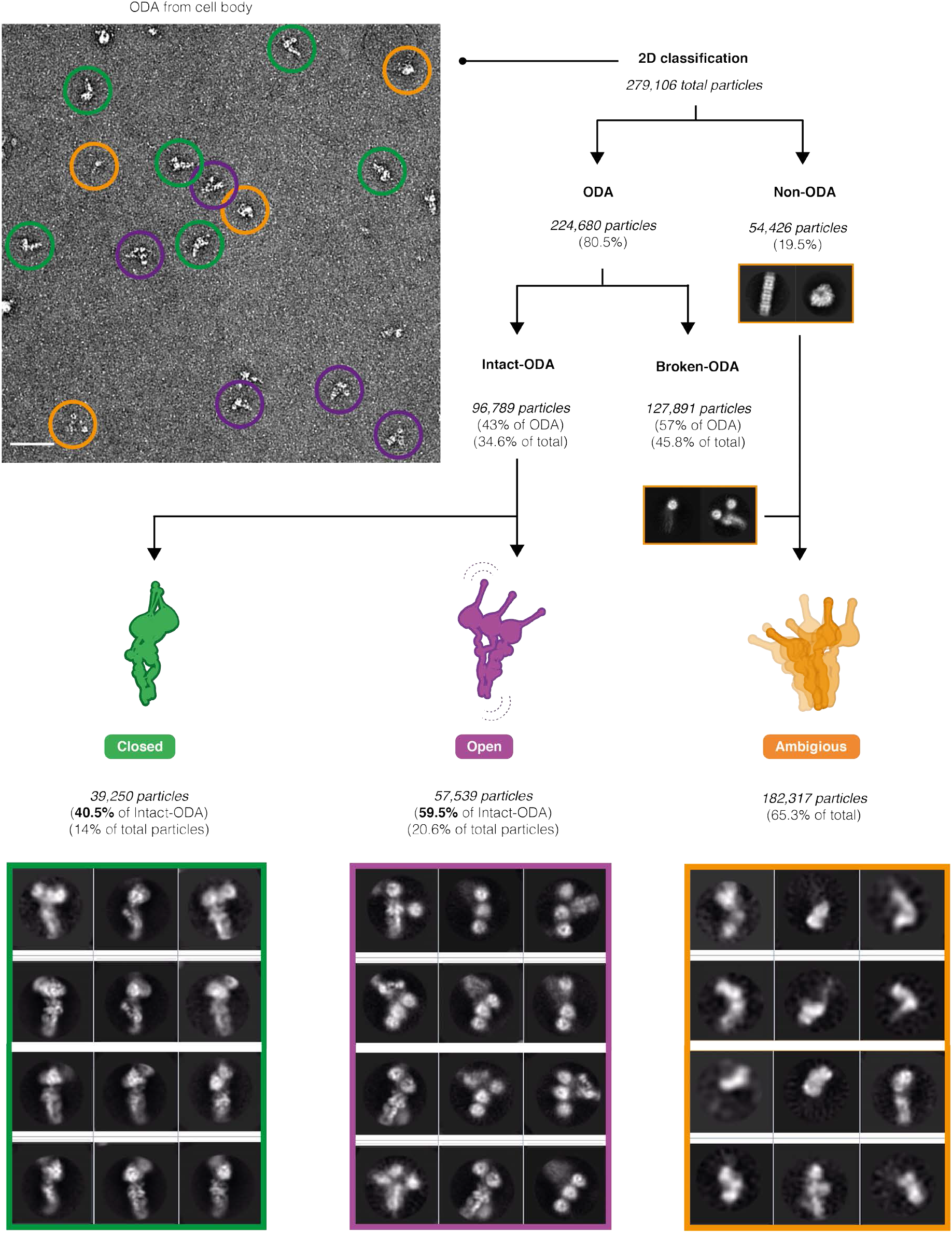
ODAs purified from cell body have a closed conformation. Representative micrograph showing negatively-stained ODA particles purified from deciliated *Tetrahymena* cell bodies. 2D classification scheme shows ∼270,000 closed ODA (green), open ODA (purple) and ambiguous particles (orange) sorted into respective class averages. Representative 2D class averages corresponding to 40% closed intact ODA particles (green), 60% open intact ODA particles (purple box).

**Figure S5.**
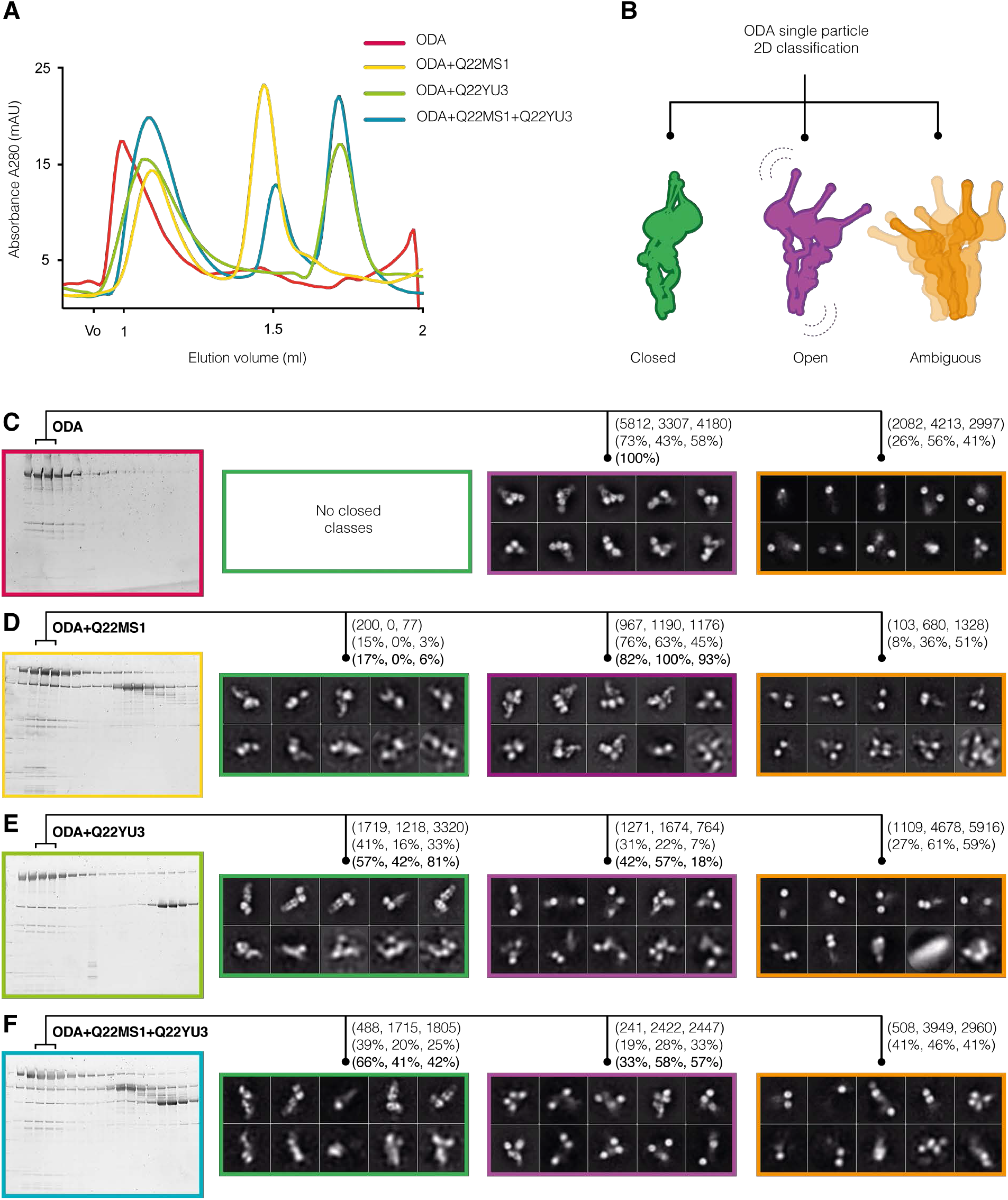
Single particle 2D classification scheme. **(A)** Representative gel filtration traces from reconstitution experiments. **(B)** 2D classification strategy used to sort ODA particles into three conformational states, closed (green), open (purple) and ambiguous (orange). **(C-F)** SDS-PAGE gel images showing fractions corresponding to traces in (A). Peak fractions were used for negative staining and particles were counted after several rounds of 2D classification. Representative 2D class averages from a final round of classification used for final subset selection and particle counting are shown. Note: ODA only sample has no closed particles.

**Figure S6.**
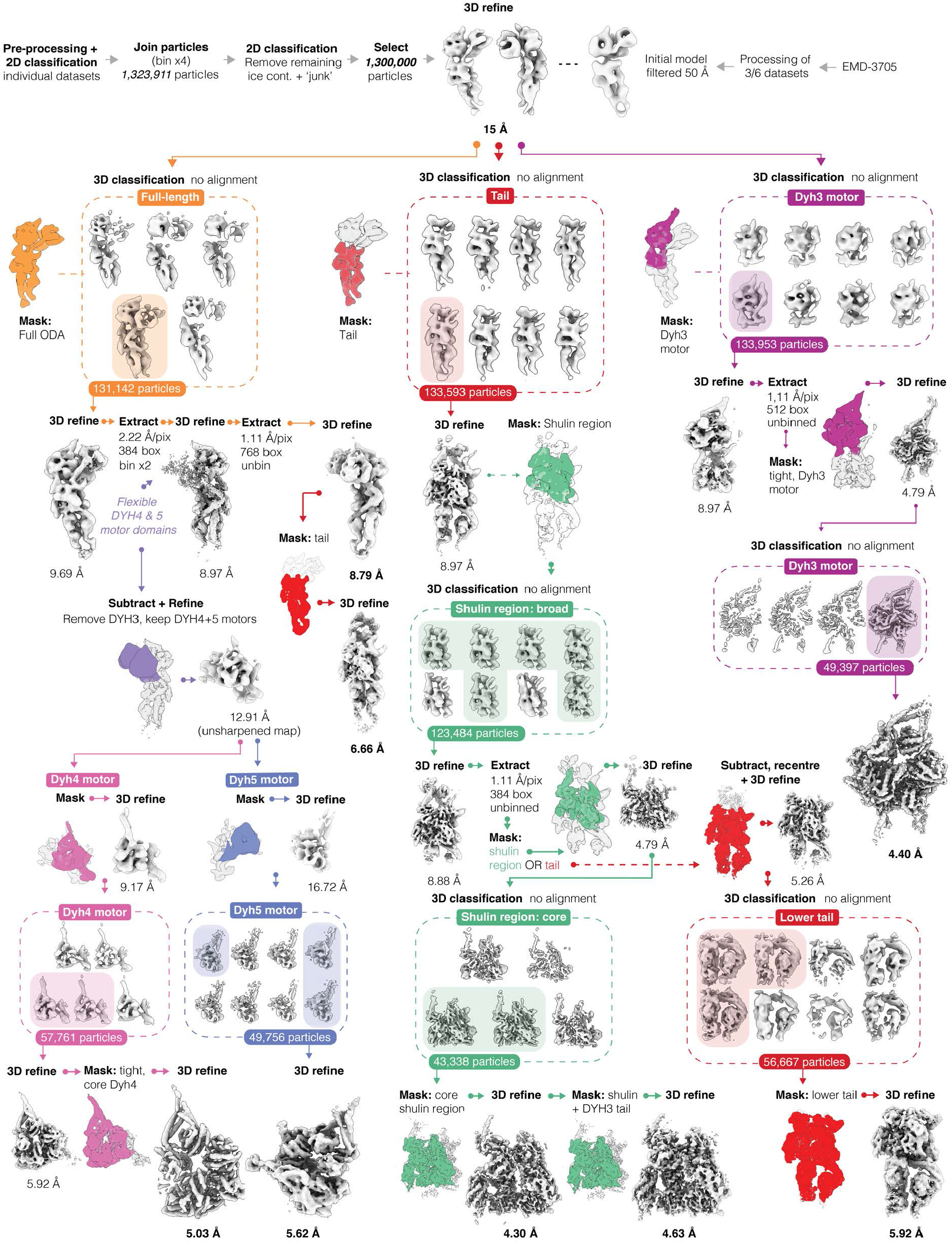
Overview of ODA image processing. A consensus 3D refinement was used as the basis for three main arms of processing in RELION-3.1: (1) Full-length ODA, Dyh3 and Dyh4 (2) Tail and Shulin region (3) Dyh3 motor.

**Figure S7.**
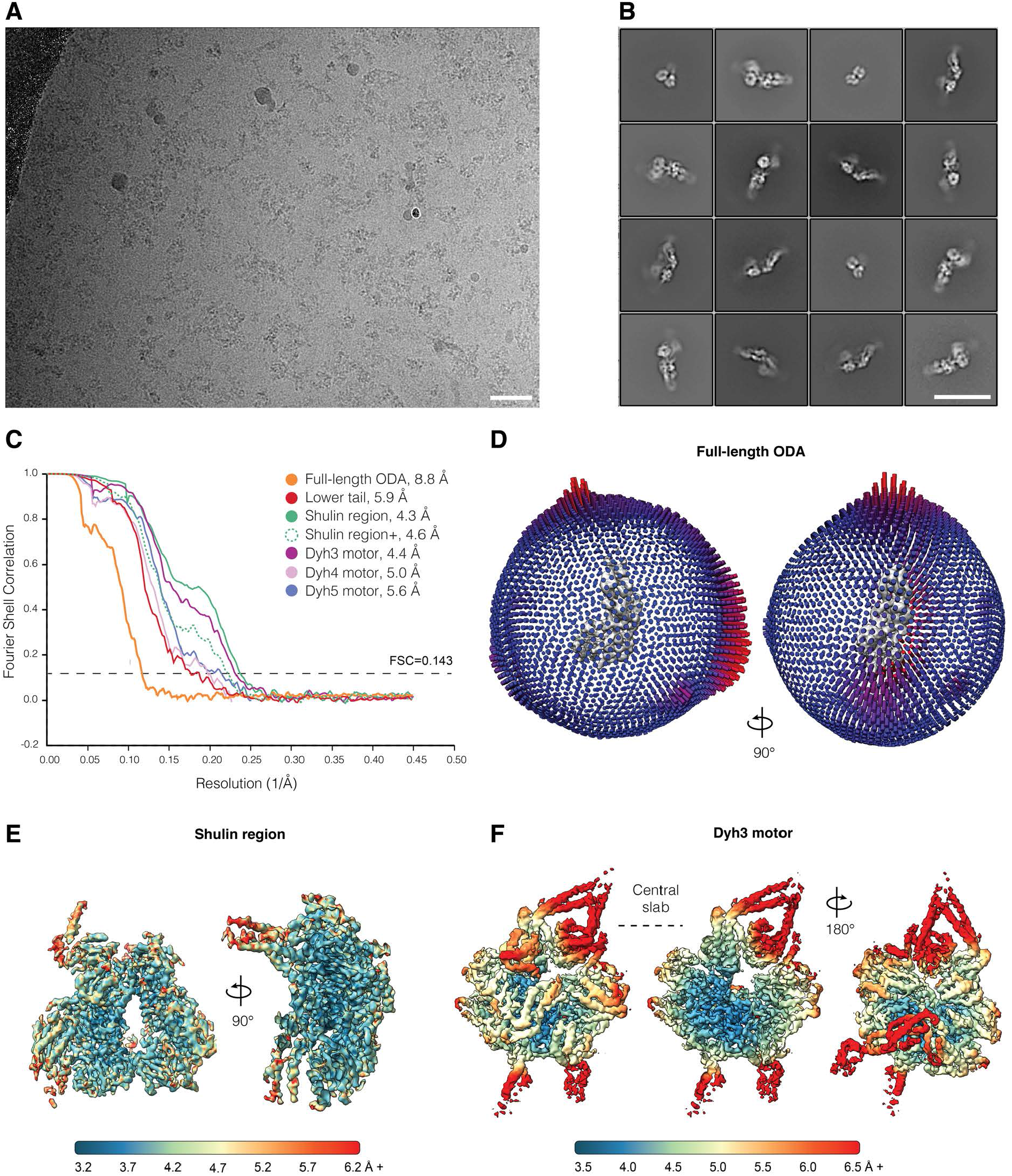
Validation of map and model quality. **(A)** Representative micrograph acquired with a K3 camera operated in counting mode. **(B)** Selection of some of the ODA-Shulin 2D class averages obtained from the combined datasets. **(C)** Gold standard Fourier shell correlation (FSC) curves for the different maps used in this study (FSC=0.143). **(D)** Angular distribution of particles contributing to the overall ODA structure. **(E-F)** Local resolution as determined by RELION-3.1’s local resolution implementation.

**Figure S8.**
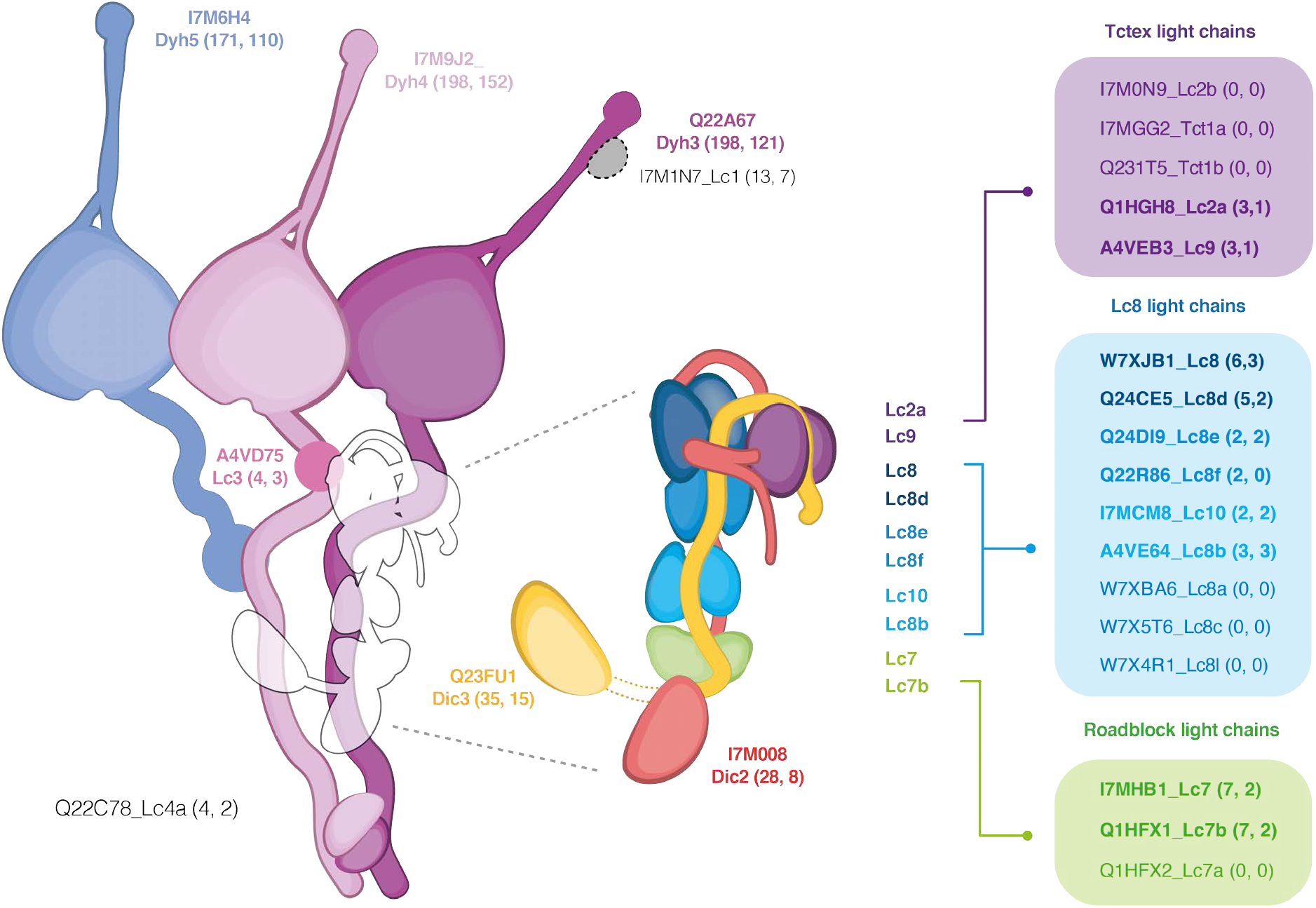
Mass spectrometry guided assignment of ODA subunits. Schematic summarizing mass spectrometry (MS) data for determination of ODA subunit composition. ODAs purified from IC3-ZZ-FLAG ciliary immunoprecipitations and isolated from wildtype cilia, comprise the subunits in bold. Peptide counts from both MS runs are shown in brackets next to protein. These data guided subunit assignment and accurate model building of the LC-tower region. For assignment, all paralogous LC sequences for Tctex-like, LC8-like and roadblock-like LC subunits (Tetrahymena Genome Database and reported previously (44)) are shown in purple, blue and green boxes respectively. LCs with best fits and confirmed via MS are in bold. Exclusive presence of Lc7, Lc7b, Lc8b and Lc10 subunits in the MS dataset allowed tentative assignment for these chains. Chains in black are not in the structure.

**Figure S9.**
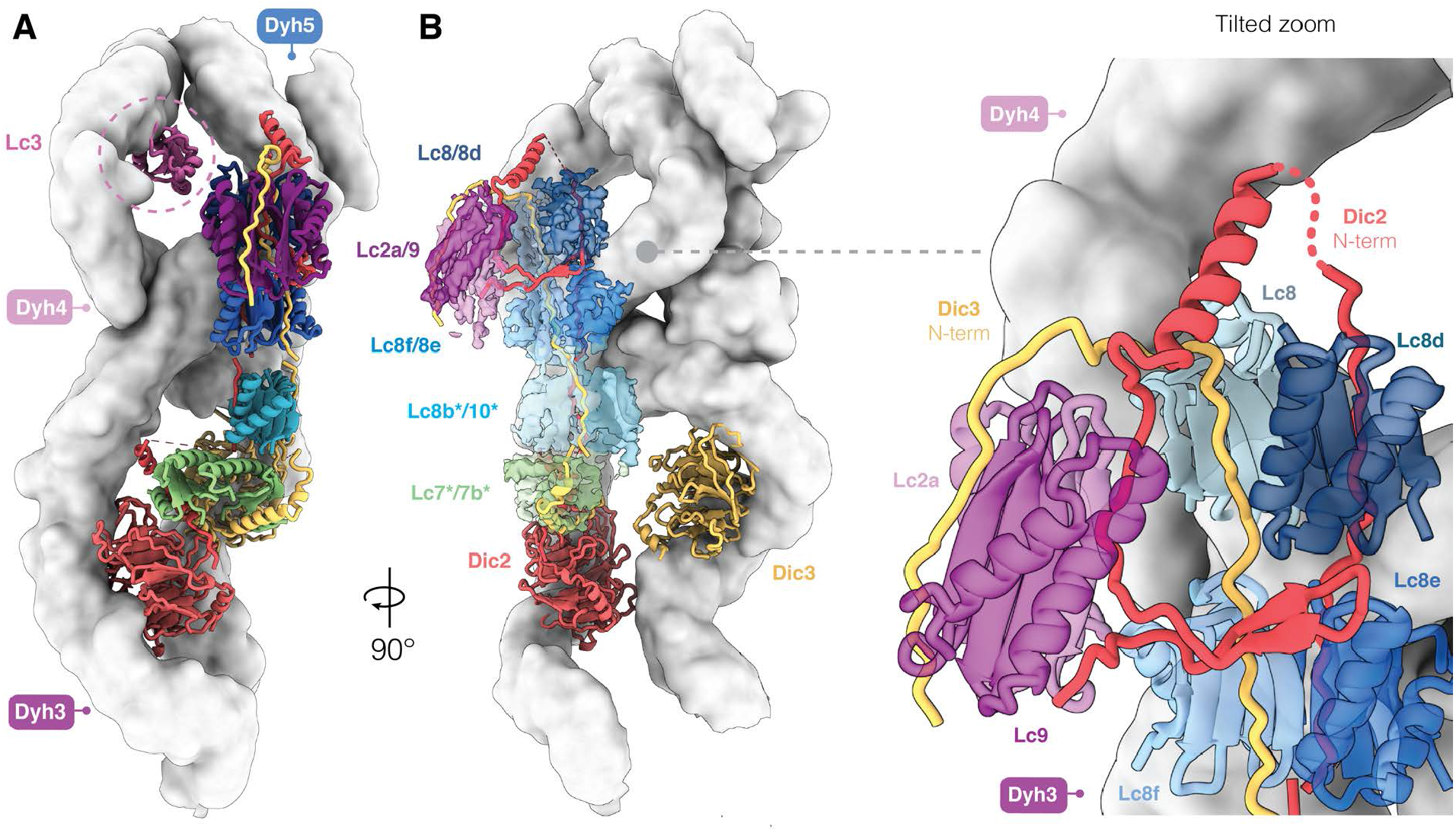
Architecture and organization of the LC tower. **(A)** The lower and upper regions of the LC tower are stacked against the Dyh3 chain whereas Lc3 contacts Dyh4 behind the main tower. **(B)** Focus on DIC N-termini and the interactions they make with the LCs. LC subunits with an * are tentatively assigned based on mass spectrometry data. The zoom inset highlights the looping of the Dic2 N-terminus around the upper LC tower. This figure depicts atomic model of this region with the heavy chains shown as filtered surface representation (white) and LCs shown as cryo-EM density in (B) and ribbon in (B, inset).

**Figure S10.**
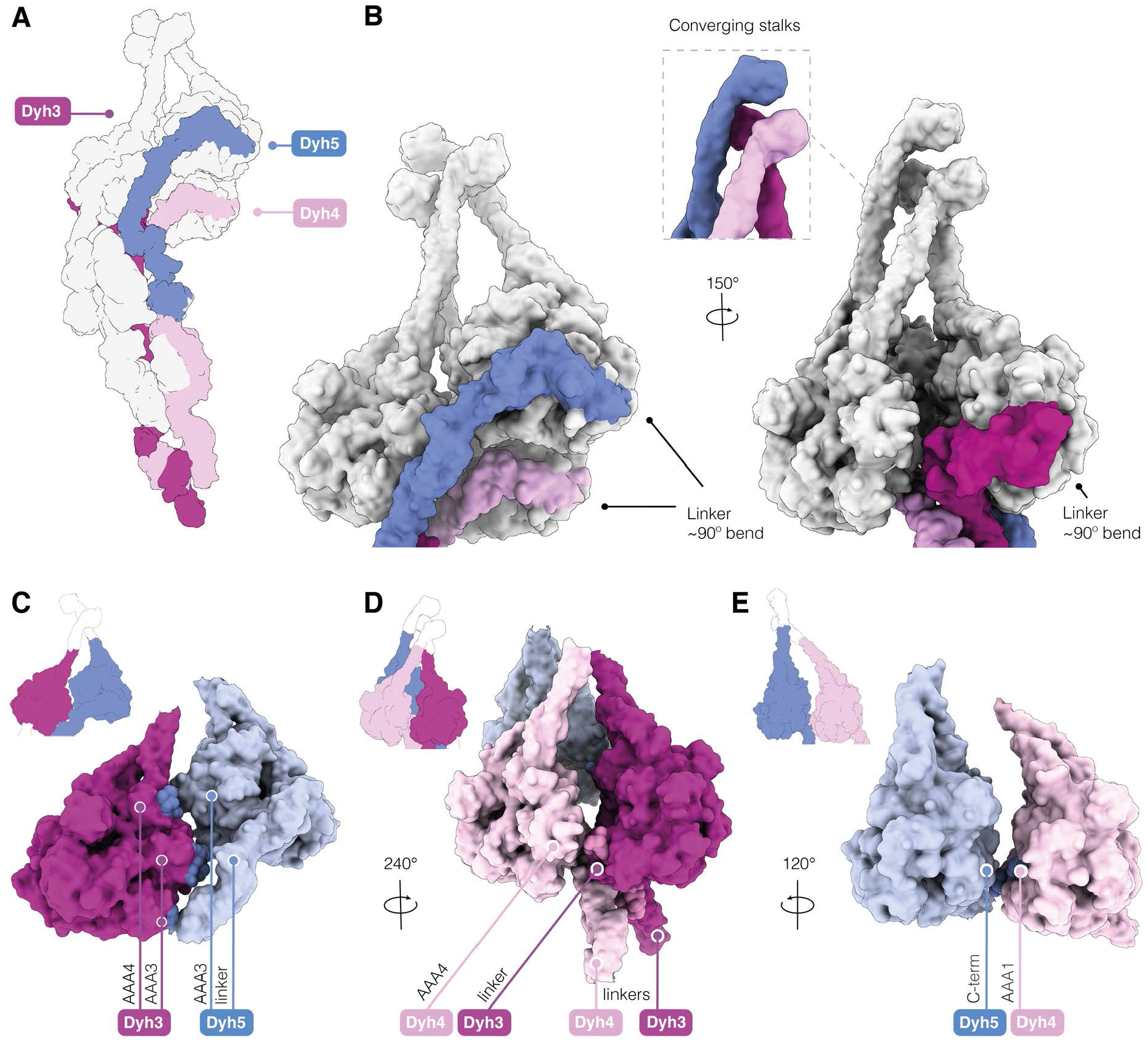
ODA motors are locked in an inactive state. **(A-B)** Shulin stabilizes the closed conformation of ODA indirectly through several contacts with the tail. Each motor domain has its force-producing linker bent at a 90° angle, indicating an inactive pre-power stroke conformation. The trajectory of the stalks suggests that they converge at their microtubule binding domains. **(C-E)** Motor-motor contacts are shown as spheres. Figure shows filtered surface representation of the atomic model.

**Figure S11.**
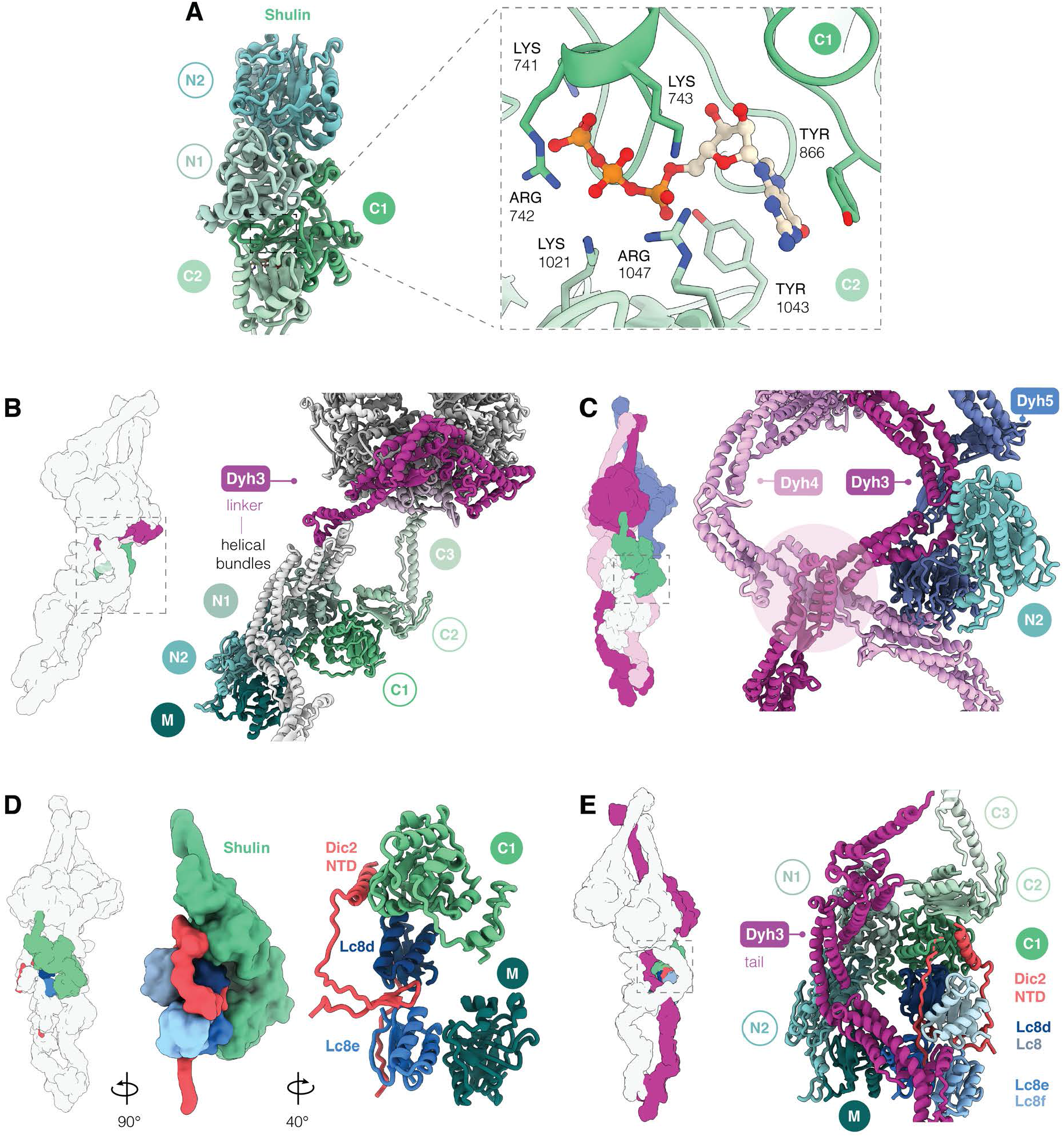
Shulin binds GTP and contacts the LC tower. **(A)** Nucleotide binds at the interface of the Shulin C1 and C2 domains. Residues in the nucleotide pocket are shown. **(B)** Shulin’s contacts with Dyh3 tail and motor, holds the linker in a pre-powerstroke conformation. **(C)** Contacts between Dyh3 and Dyh4 tail helical bundles, stabilized by Shulin, are shown. **(D)** Shulin contacts with Lc8e, Lc8d and Dic2 are shown. Dic2 N-terminal domain (NTD) loop wedged between the two LC subunits is shown. **(E)** Shulin bridges Dyh3 and LC tower, packing the latter against the Dyh3 tail. Shulin domains making contacts with ODA subunits are in filled circles.

**Figure S12.**
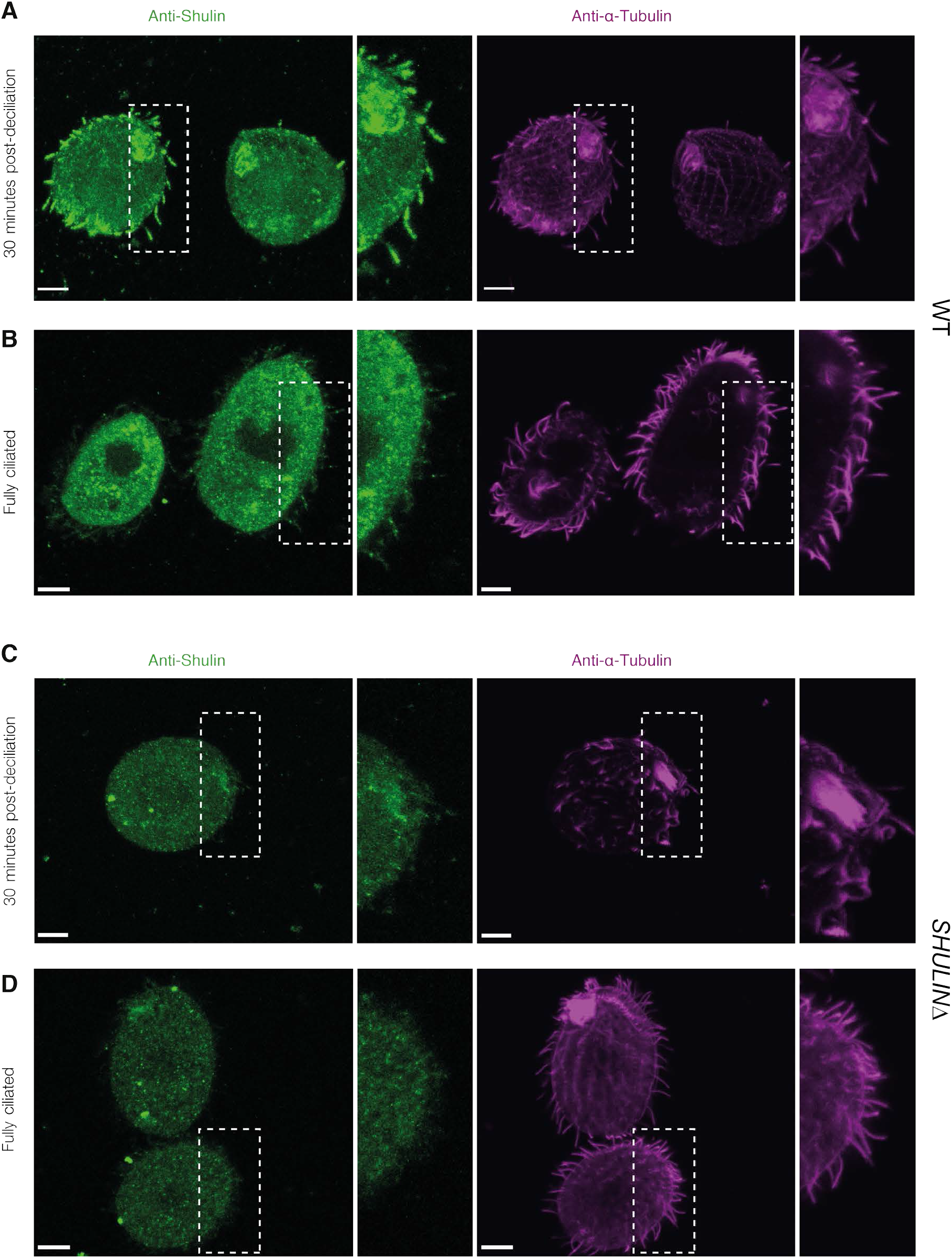
Shulin delivers ODAs during *de novo* ciliogenesis. **(A)** Immunostaining of cells regenerating motile cilia after deciliation shows Shulin enters actively extending cilia marked by acetylated α-tubulin. Insets highlight ciliary staining for Shulin. **(B)** Immunostaining of cells with fully assembled cilia shows Shulin localizes to the cytoplasm. Insets highlight residual ciliary staining for Shulin. **(C-D)** *SHULIN*Δ cells stained as above, lacking Shulin signal serve as immunostaining controls. Note: differences in tubulin staining between regenerating cilia and fully assembled cilia reflect differences in tubulin acetylation which occurs after microtubule assembly in *Tetrahymena* axonemes (45).

**Table S1.** Quantitative mass spectrometry hits of novel ODA cell body interactors. Label-free quantitative MS list of high abundance proteins specifically co-precipitating with IC3-ZZ-FLAG from quadruplicate runs. Shulin and Q22MS1 are highlighted. Proteins in bold letters are part of the preassembled ODA complex in the cell body. MW = molecular weight, FC = log_2_ fold-change, P-value significance is in -log_10_ scale.

| UNIPROT ID | PROTEIN ID | MW (kDa) | P-value | FC |
| --- | --- | --- | --- | --- |
| **I7MHB1_TETTS** | **Lc7** | **15** | **0.53** | **4.85** |
| **I7M9J2_TETTS** | **Dyh4/ODA β-HC** | **530** | **1.82** | **4.57** |
| **Q22A67_TETTS** | **Dyh3/ODA γ-HC** | **534** | **2.23** | **4.51** |
| **Q22MS1_TETTS** | **NDK-domain Protein** | **222** | **0.58** | **4.26** |
| **Q22YU3_TETTS** | **Shulin** | **140** | **1.03** | **4.12** |
| I7LX00_TETTS | Haemoglobin-like Protein | 14 | 0.70 | 4.03 |
| **I7M1N7_TETTS** | **Lc1** | **23** | **1.55** | **4.03** |
| **A4VD75_TETTS** | **Lc3** | **14** | **0.62** | **4.01** |
| Q231E4_TETTS | Ttp95-like Protein | 126 | 1.67 | 3.88 |
| **I7M008_TETTS** | **Dic2/ ODA IC** | **78** | **1.27** | **3.87** |
| **Q1HGH8_TETTH** | **Lc2a** | **16** | **0.46** | **3.63** |
| **I7M6H4_TETTS** | **Dyh5/ODA α-HC** | **475** | **2.98** | **3.54** |
| **Q23FU1_TETTS** | **Dic3/ ODA IC (Bait)** | **77** | **4.55** | **3.52** |
| W7X3J8_TETTS | Na/Ca Exchanger Protein | 52 | 1.06 | 3.51 |
| **Q1HFX4_TETTH** | **Lc4a** | **18** | **0.46** | **3.42** |
| Q23QY3_TETTS | Uba1 | 117 | 0.46 | 3.40 |
| Q22AU5_TETTS | SLEI-family Protein | 283 | 0.44 | 3.34 |
| I7M9Z0_TETTS | Copg2 | 283 | 0.46 | 3.33 |
| **W7XJB1_TETTS** | **Lc8** | **13** | **0.46** | **3.32** |
| I7MFH3_TETTS | Cytochrome C-like Protein | 12 | 0.46 | 3.31 |
| Q23AC8_TETTS | Dnajb6 | 30 | 0.60 | 3.28 |
| Q24FD6_TETTS | Dnaja2 | 47 | 1.32 | 3.24 |
| W7XEU3_TETTS | Cystatin-domain Protein | 10 | 0.46 | 3.14 |

**Table S2.** Cryo-EM data collection and processing

|  | ODA-S_1 | ODA-S_2 |
| --- | --- | --- |
| **Data collection and processing** |  |  |
| Microscope | TFS Titan Krios<br>(LMB) | TFS Titan Krios (eBIC) |
| Detector | K3 | K3 |
| Magnification (nominal) | 81,000x | 81,000x |
| Pixel size (Å) | 1.11 | 0.53 (super- res) |
| Energy filter slit width (eV) | 20 | 20 |
| Voltage (kV) | 300 | 300 |
| Number of frames | 66 | 54 |
| Exposure (s) | 2.6 | 3 |
| Voltage (kV) | 300 | 300 |
| Electron exposure on sample (e-/Å<sup>2</sup>) | 52 | 52 |
| Target defocus range (µm) | 1.2-3 | 1.5-3 |
| Number of sessions | 6 | 1 |
| Number of collected movies | 40,158 | 10,463 |
| Particle numbers after removing graphene oxide<br>artifacts/ice contamination |  | 1,300,000 |
| Final particle numbers and map resolution at<br>FSC=0.143 (Å) | Full-length: 131,142; **8.8** (EMD-11576)<br>Tail: 131,142; **6.7** (EMD-11577)<br>Ordered tail: 56,667; **5.9** (EMD-11578)<br>Shulin region: 43,338; **4.3** (EMD-11579)<br>Extended Shulin region: 43,338; **4.6** (EMD-11580)<br>Dyh3 motor region: 49,397; **4.4** (EMD-11581)<br>Dyh4 motor: 57,761; **5.0** (EMD-11582)<br>Dyh5 motor: 49,756; **5.6** (EMD-11583)<br>Extended Dyh5 motor: 49,756; **9.3** (EMD-11584) |  |

**Table S3.**
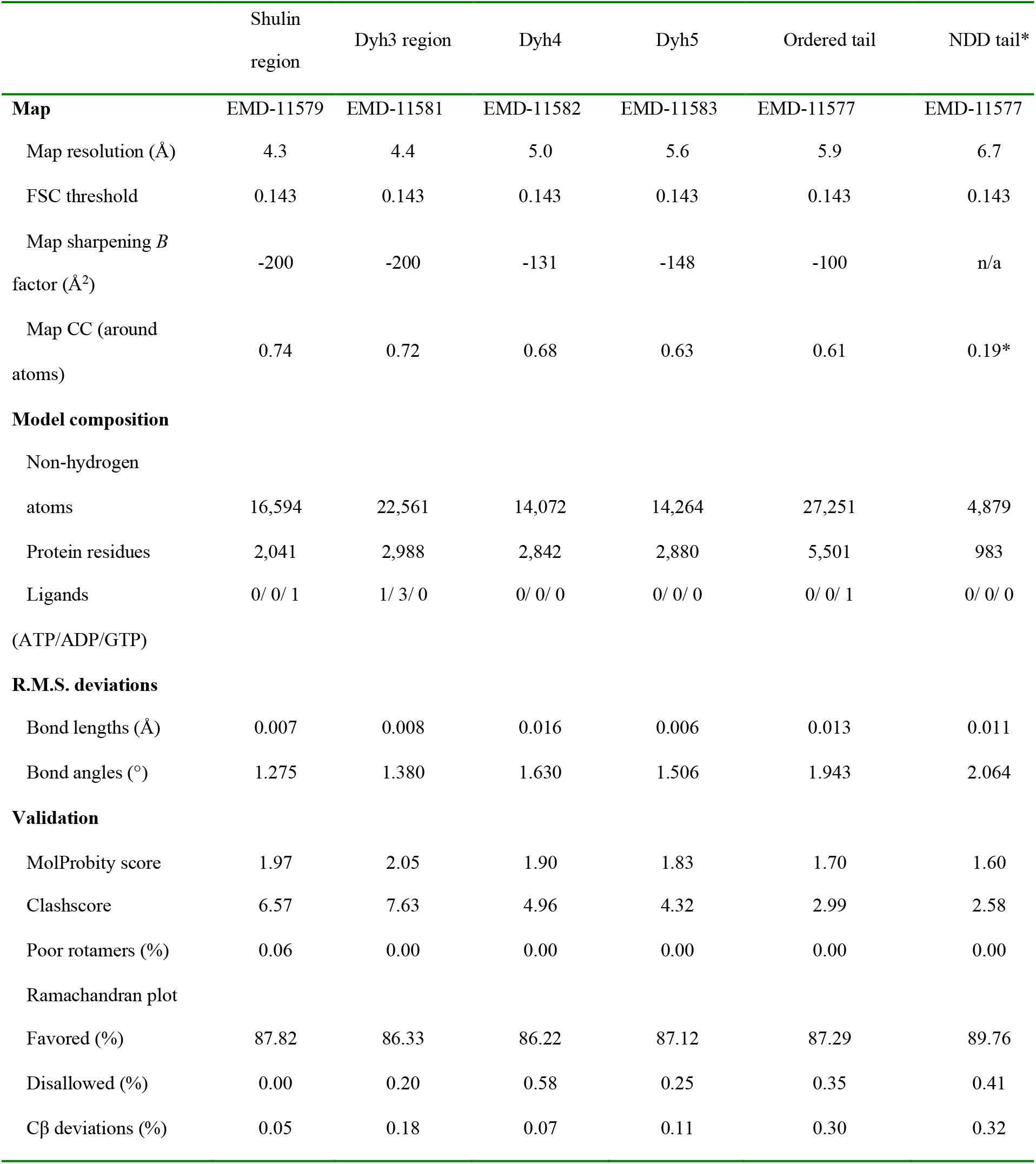
Refinement and validation statistics. * The flexible NDD region of the tail was modeled into low resolution density at low threshold.

**Movie. S1**. Movie shows slow moving cilia (arrows) in Q22YU3 and Q22MS1 mutant cells compared to the wildtype cells which have fast-moving cilia (arrows).

**Movie. S2**. Movie shows slow moving cilia (arrows) in the temperature sensitive OAD1 C11 mutant cell, which has reduced ODAs in cilia compared to the wildtype cell which has normal ciliary ODA levels and fast-moving cilia (arrows).

**Movie. S3**. Movie depicting the model of ODA bound by Shulin (green). Subunits are listed and contacts made by Shulin with the various ODA subunits are shown as green spheres.

